# A conserved fungal transcription factor domain drives protein-protein interactions and necrotrophic effector-mediated virulence in *Parastagonospora nodorum*

**DOI:** 10.64898/2025.12.09.693112

**Authors:** Shota Morikawa, Carlia Herbst, Evan John, Daniel Croll, Carl Mousley, Bernadette Henares, Kar-Chun Tan, Callum Verdonk

## Abstract

The fungal pathogen *Parastagonospora nodorum* utilises necrotrophic effectors (NEs) to cause chlorosis and necrosis on wheat. NE expression is mediated by an assortment of transcription factors (TF), the most well-characterised of which is PnPf2. Orthologues of PnPf2 regulate virulence in phytopathogenic fungi across the Ascomycete fungal lineage, yet their protein architecture remains functionally uncharacterised. These orthologues are characterised by the archetypal N-terminal Zn_2_Cys_6_ zinc-finger DNA-binding domain (DBD), a conserved yet poorly characterised middle-homology region (MHR) and a C-terminal disordered region. We investigated the role of each of the three domains through PnPf2 truncation mutants *in situ*. This revealed the conserved MHR of PnPf2 is required for the development of disease symptoms on wheat, but also that the C-terminal disordered region in-part modulates NE expression. Domain-interaction analysis through yeast-2-hybrid (Y2H) screening reveals PnPf2 forms a homodimer mediated by the MHR, indicating its importance mediating protein-protein interactions. Using the MHR as a bait in library-scale protein-protein interaction Y2H assays, we identified the COP9-signalosome protein PnCsn6 as a key interaction partner. PnCsn6 is essential for disease symptoms during *P. nodorum* infection on wheat, including NE expression. Our study presents the first domain-level functional investigation this virulence-regulating TF orthologue, the results of which underpin the essential role of the MHR in driving protein-protein interactions and effector regulation. It also reveals an essential protein-signalling pathway with which PnPf2 directly interacts and shares regulatory control of virulence.

## INTRODUCTION

Phytopathogenic fungi utilise small, secreted effector molecules that modulate the outcome of a plant-microbe interaction (Friesen et al., 2008, Kanyuka et al., 2022, McDonald and Solomon, 2018, Lo Presti et al., 2015). These include fungal pathogens that employ necrotrophic effectors to disable a host ahead of infection by interacting with a compatible dominant susceptibility gene. A compatible interaction results in host tissue chlorosis and/or necrosis, allowing the pathogen to infect (McDonald and Solomon, 2018, Lo Presti et al., 2015, Kariyawasam et al., 2023). Effector expression and activity is intricately regulated at multiple levels through epistasis, chromatin remodelling, sub-cellular localisation, gene copy-number variation and transcription factors (TFs) (Tan and Oliver, 2017, Amaike et al., 2008, Aboukhaddour et al., 2012). TFs are DNA-binding regulatory proteins that control various functions associated with fungal pathogen development, such as vegetative growth, secondary metabolism and virulence factor expression (Shelest, 2017, John et al., 2021, Tan and Oliver, 2017). TFs carry specialised DNA-binding domains (DBDs), often at their N-termini, which facilitate gene expression (John et al., 2022, Shelest, 2008). A major family of TFs that are unique to fungi carry a highly conserved middle homology region (MHR) fused to an N-terminal DBD, typically a zinc-finger of the Zn_2_Cys_6_ class. In addition to being unique, these TFs are ubiquitous among fungi and represent the largest TF family (Mayer et al., 2023, Todd et al., 2014, Tianqiao et al., 2021). Despite this, the role of the MHR is understudied, particularly in plant pathogenic fungi. In model fungi, it has been shown that the MHR is composed of eight consecutive conserved α-helical motifs with broadly defined roles often associated with nuclear receptors or DNA-complex assembly (Näär and Thakur, 2009, Poch, 1997, Mayer et al., 2023). A well-characterised example of a Zn_2_Cys_6_ DBD-MHR TF is the galactose-induced Gal4 from baker’s yeast *Saccharomyces cerevisiae* (Traven et al., 2006). The activation of Gal4 is dictated by a nine-amino acid (aa) transactivation domain (9aaTAD) in the C-terminus of the protein (Piskacek et al., 2007). Despite this, the function of the MHR in Gal4 has not been fully elucidated.

In the wheat fungal pathogen *Parastagonospora nodorum*, the TF PnPf2 has a Zn_2_Cys_6_ DBD-MHR architecture (Rybak et al., 2017, John et al., 2024). PnPf2 was originally discovered as an orthologue of AbPf2 in *Alternaria brassicicola*, a fungal pathogen of *Brassica* spp. (Cho et al., 2013). The deletion of *PnPf2* and *AbPf2* in their respective pathogens results in the abolishment of virulence but has minimal impact on vegetative growth, suggesting that Pf2 orthologues are essential for full virulence but are not major contributors to fungal development (Cho et al., 2013, Rybak et al., 2017). PnPf2 is a positive regulator of the necrotrophic effector (NE) genes *SnToxA* and *SnTox3* (Rybak et al., 2017, Jones et al., 2019) and has a broader role in regulating carbohydrate-active enzymes and nutrient assimilators (John et al., 2024). LmPf2 from the hemibiotroph *Leptosphaeria maculans* acts antagonistically with the chromatin remodeler KMT1 for direct effector regulation (Clairet et al., 2024), implicating the role of Pf2 in complex genomic reassembly for its regulation. All characterised Pf2 orthologues are conserved within the Ascomycota phylum and harbour a modular domain architecture typical of Zn_2_Cys_6_ TFs: with an N-terminal DBD, a MHR and a low-homology disordered C-terminus (Clairet et al., 2024, Cho et al., 2013, See and Moffat, 2023, John et al., 2024, Habig et al., 2020, Oh et al., 2016).

Despite the abundance of MHR in fungal TFs (Shelest, 2008, Todd et al., 2014, Mayer et al., 2023), little is known about the function of the MHR domain within transcription factors. Current studies limit their investigations to entire gene disruptions containing the MHR (John et al., 2021), restricting the discovery of domain-specific roles within TFs. Furthermore, little is known about the direct relationship between MHR function and its role in virulence progression within phytopathogenic fungi, enabling a new avenue to pursue to address plant-pathogen paradigms. Here, we elucidated the functions of each PnPf2 region, including refined characterisation of the conserved and ubiquitous MHR, and their association with the pathogenicity of *P. nodorum* through gene truncation-mutant investigations. Utilising a yeast-2-hybrid (Y2H) library-screening strategy, we uncovered a number of interacting partners of PnPf2. Through deeper investigation, we reveal a key regulatory role of the PnPf2-interacting COP9-signalosome subunit PnCsn6 in fungal development, asexual sporulation and virulence.

## RESULTS

### An analysis of Pf2 domain conservation

PnPf2 is a transcriptional regulator of the NE genes *SnToxA* and *SnTox3* and carries a Zn_2_Cys_6_ DBD-MHR domain architecture typical of fungal TFs (John et al., 2022, Rybak et al., 2017). Previous analysis of Pf2 proteins across fungal orthologues indicated relatively high conservation across both the N-terminal Zn_2_Cys_6_ domain (*Zn*) and the central MHR, with poor conservation at the C-termini predicted disordered region (*Dis*) (John et al., 2024) (**Supplemental Table S1**). Given the low conservation of the C-terminal region amongst Pf2 orthologues between fungi especially at the disordered C-terminal region, we sought to uncover whether this low-conservation exists in PnPf2 between strains within *P. nodorum*. An aa sequence alignment of PnPf2 from a diverse global collection of 161 isolates (Pereira et al., 2021) - collected from Australian, European, North and South American, and Middle Eastern wheat fields - showed that the PnPf2 protein is always present, and has a high conservation > 99% overall aa identity including the C-terminal disordered region (**Table 1**). This suggests that each domain individually may play an important role for PnPf2 function within *P. nodorum*. Furthermore, a DNA sequence alignment of the 161 isolate *PnPf2* upstream sequences showed a 100% sequence conservation within a 447-bp upstream region containing the promoter of *PnPf2*, indicating PnPf2 regulation is well-conserved between isolates (**Supplemental Figure S1**).

**Table 1:** The aa sequence conservation of PnPf2 and Pf2 orthologues shows high conservation within a global *P. nodorum* isolate panel but low conservation between species. The aa residue positions are based on PnPf2 from *P. nodorum* SN15. “Identity” refers to the percentage of identical aa residues at a given position in the alignment, while “positive” refers to the percentage of similar aa residues at a given position. *Zn* is the putative Zn_2_Cys_6_ DBD, MHR is the middle homology region and *Dis* is the disordered region.

| Pf2 alignment | *Zn* (≤63 aa) identity [positive] | MHR (64-497 aa) identity [positive] | *Dis* (≥498 aa) identity [positive] |
| --- | --- | --- | --- |
| 161 *P. nodorum* isolates | 99.96% [100%] | 99.9% [99.9%] | 99.5% [99.8%] |
| 12 fungal species* | 67.3% [77.3%] | 38.6% [56.8%] | 12.9% [19.6%] |

\*The 12 fungal species from the orders: Pleosporales, Capnodiales, Helotiales, Magnaporthales, Hypocreales and Sordariales are based on the phylogenetic analysis described in John et al. (John et al., 2024)(**Supplemental Table S1**).

### PnPf2 disordered region is dispensable for virulence but needed for full effector regulation

The high conservation of Pf2 homologues between isolates is particularly interesting, given the diverse effector profile within variable strains of *P. nodorum* (McDonald et al., 2013). This suggests that there must be a conserved functional role for each domain within Pf2 (Jones et al., 2019). To determine the function of each PnPf2 domain in effector regulation and virulence, we developed a progressive-series of *PnPf2* truncations *in situ* made according to the domain region-boundaries outlined in John et al. (John et al., 2024)(**Figure 1A**; **Table 2**). With these *PnPf2* gene-truncation mutants, we performed phenotypic assays assessing vegetative morphology and infection on the highly susceptible wheat cultivar (cv.) Halberd (*Tsn1*, *Snn1*, *Snn3*). The full-length *PnPf2*-mutant *PnPf2_1-653_*, which carried an intact *PnPf2* gene and selectable transformation marker, exhibited identical lesion sizes and vegetative morphology as the SN15 wildtype (**Figure 1B**); indicating the resistance gene has no effect on observable phenotypes. C-terminal partial-truncations of the predicted *Dis* domain *PnPf2_1-539_* and *PnPf2_1-529_* were also comparable in disease morphology to SN15 in detached leaf assays (DLA), suggesting the putative 9aaTAD present within the PnPf2 *Dis* domain was not required for observable virulence in wheat. Concordantly, the PnPf2 *Dis*-truncation gene *PnPf2_1-497_,* which retains the complete *Zn*-MHR, exhibited disease symptoms on wheat comparable to SN15 (**Figure 1C**). In contrast, the MHR-*Dis* truncated PnPf2 variant *PnPf2_1-63_,* which contained only the Zn_2_Cys_6_ putative DBD *Zn*, was unable to induce disease and was comparable to the complete *PnPf2-*deletion strain *pnpf2ko*. Taken together, we suggest the MHR is important for PnPf2-mediated disease progression and chlorotic/necrotic lesion development during wheat infection.

**Figure 1:**
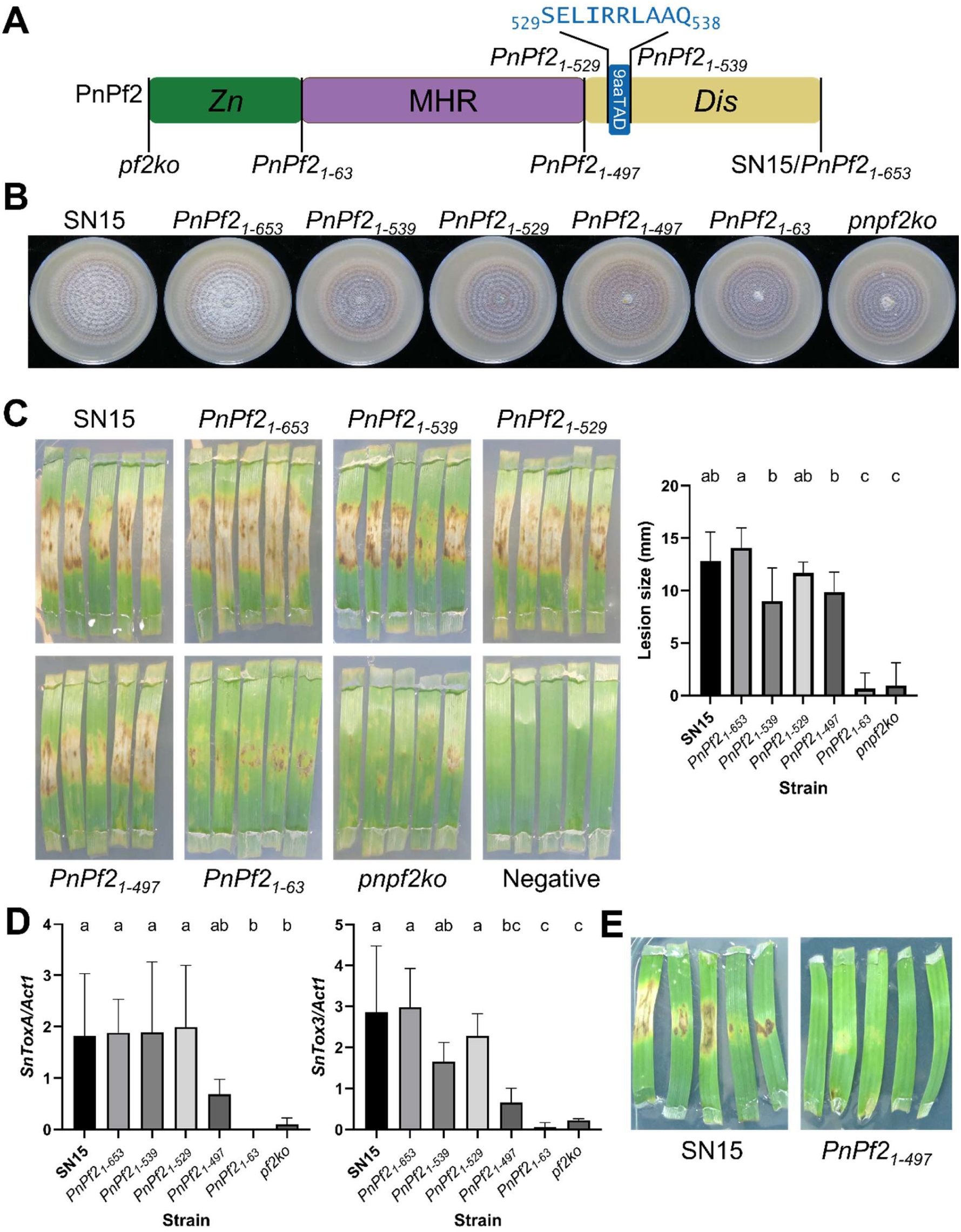
PnPf2 middle homology region (MHR) is required for virulence of *P. nodorum* on wheat. (**A**) A schematic showing the various PnPf2 truncation mutants. *Zn* is the putative Zn_2_Cys_6_ DNA-binding domain (aa 1-63), MHR is the middle homology region (aa 64-497), 9aaTAD is the predicted nine amino acid transactivation domain (aa 530-539) and *Dis* is the poorly conserved disordered region (aa 498-653). (**B**) Vegetative growth of PnPf2-truncated mutants showing minimal growth abnormalities compared to SN15 and closely matched with the phenotype observed previously (Rybak et al., 2017). (**C**) Plug detached leaf assay (DLA) of PnPf2 truncation mutants 10 days post inoculation (cv. Halberd) with an agar plug. Negative indicates the Tween20-only treatment. The plot shows the lesion sizes of each PnPf2 truncation mutant with letters above the bars representing statistical significance according to a one-way ANOVA and Tukey’s HSD (n = 5; *p* < 0.05). **(D)** PnPf2 truncations resulted in dysregulation of necrotrophic effector (NE) gene expression. Quantitative PCR of NE genes (*SnToxA* and *SnTox3*) normalised to the housekeeping actin (*Act1*) during infection of wheat (cv. Halberd) three days post inoculation where NE expression is maximal. Different letters above the bars indicate significant difference between strains within a plot according to a one-way ANOVA and Tukey’s HSD test (n = 3; *p* < 0.05). **(E)** Plug DLA of SN15 compared with the C-terminal *Dis* truncation strain PnPf2_1-497_ on the SnTox3-sensitive cv. Wyalkatchem 10 days post inoculation. All other PnPf2-truncation strains DLAs are available in **Supplemental Figure S2.**

**Table 2:**
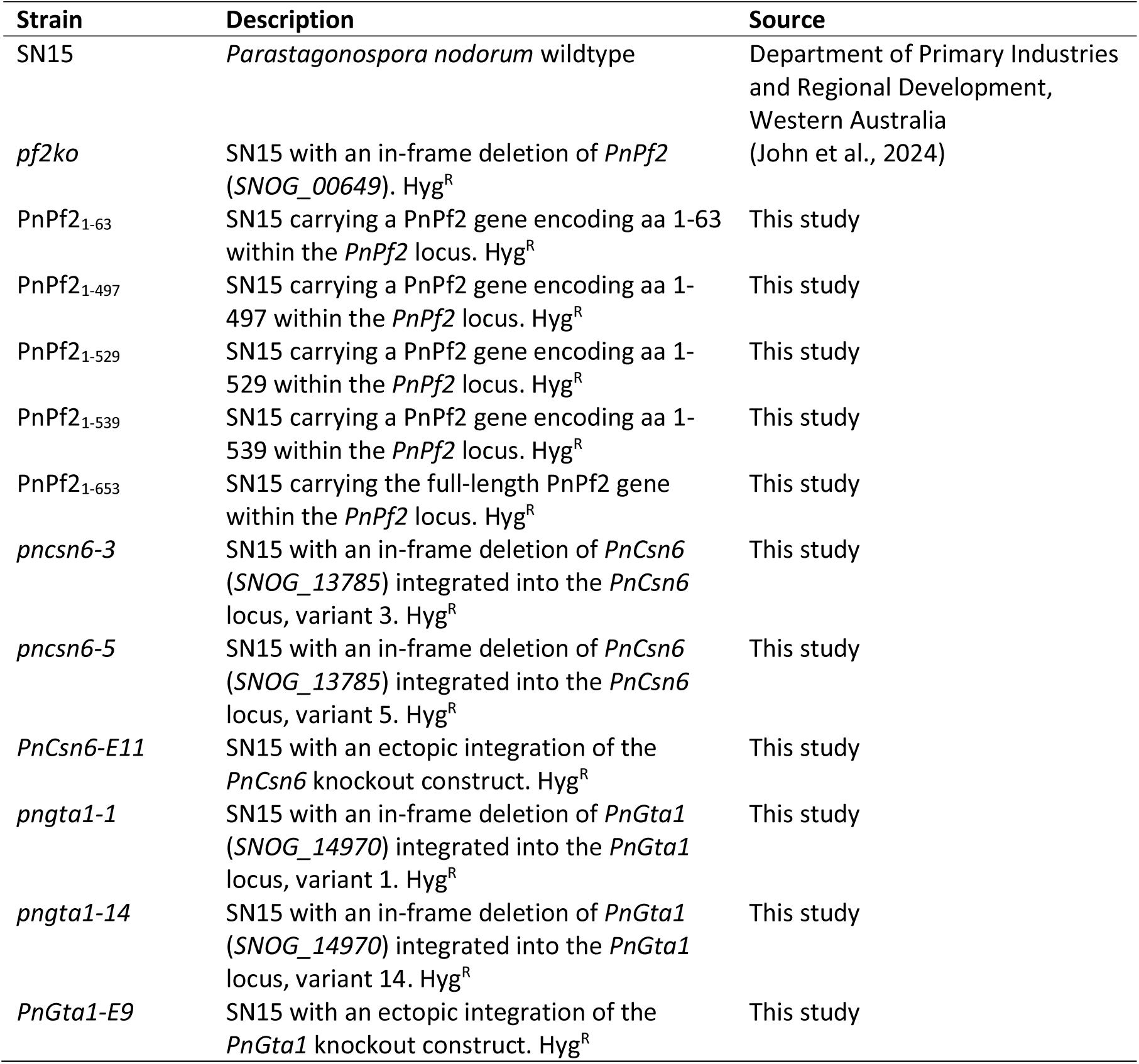
Fungal strains used throughout this study.

Given the impact of PnPf2 truncations on lesion development during host infection, we investigated how each truncation influenced the relative expression of the PnPf2 mediated effectors *SnToxA* and *SnTox3*. *PnPf2_1-653_*, along with the *Dis*-truncated variants *PnPf2_1-539_* and *PnPf2_1-529_*, had no significant impact on either *SnToxA* or *SnTox3* expression during *in-planta* infection compared with SN15, as determined by qRT-PCR (**Figure 1D**). A 3-to-4-fold reduction in *SnToxA* and *SnTox3* expression was observed when the entire *Dis* region was deleted in the *PnPf2_1-497_* truncation mutant, leaving only the *Zn*-MHR domains. In contrast, no expression of *SnToxA or SnTox3* was detected in the *Zn* only mutant *PnPf2_1-63_*, or in the full *PnPf2* deletion mutant *pnpf2ko*. Next, we tested whether this reduced *SnTox3* expression in *PnPf2_1-497_* was still able to induce necrosis on the SnTox3-sensitive wheat cv. Wyalkatchem (*tsn1*, *snn1, Snn3*), which does not show chlorotic or necrotic symptoms when infected with *pf2ko* (Rybak et al., 2017). Unlike SN15, no necrosis is observed for the entire *PnPf2* C-terminal *Dis* deletion strain *PnPf2_1-497_* (**Figure 1E; Supplemental Figure S2**), suggesting that the reduction of *SnTox3* expression is enough to ablate observable disease progression in DLAs on the *Snn3-*specific wheat cv. Wyalkatchem. Consistent with the disease phenotype described above, these findings show the importance of the PnPf2 MHR for NE expression and PnPf2 virulence. Furthermore, given the pronounced reduction in *SnToxA* and *SnTox3* expression between *PnPf2_1-529_* and *PnPf2_1-497_* - we conclude that the region between residues 497-529, encompassing the putative 9aaTAD and partial *Dis* region, is required for the normal expression of *SnToxA* and *SnTox3* during *in-planta* infection.

### PnPf2 MHR drives homodimerisation in yeast-2-hybrid

Zinc-finger MHR-containing proteins have been shown to homodimerise (John et al., 2021), so we next sought to investigate the potential for PnPf2 to homodimerise. Given the importance of the PnPf2 MHR for virulence and NE expression, we employed a targeted domain PnPf2-PnPf2 yeast-2-hybrid (Y2H) approach to uncover the putative dimerisation domain driving protein-protein interactions. Using the 653 aa full-length PnPf2-encoding protein as prey, we sequentially tested PnPf2 against all domains, both individually and as pairs, of PnPf2 baits. We observed that full-length PnPf2-encoding prey, ^PREY^PnPf2_FULL_, interacts with the MHR of PnPf2 through the blue colony abundance at high dilutions on the selective Y2H media (**Figure 2A**). Intriguingly, the interaction is lost - as no yeast growth was observed - with the addition of the Zn_2_Cys_6_ putative DNA-binding domain both as a C-terminal truncation PnPf2_Zn+MHR_ and as full-length PnPf2_FULL_ (**Supplemental Figure S3**). We also observed that the products of the PnPf2 MHR+*Dis* containing bait vector ^BAIT^PnPf2_MHR+Dis_ interacted with the full-length PnPf2 prey ^PREY^PnPf2_FULL_. We suspect this is indicative of endogenous yeast autoactivation, as we additionally observed growth of ^BAIT^PnPf2_MHR+Dis_ (but not ^BAIT^PnPf2_MHR_) colonies when tested against our negative control prey pGADT7 empty vector (EV) ^PREY^EV (**Figure 2A**). In summary, only the PnPf2 MHR interacts with full-length PnPf2 in Y2H without autoactivation, suggesting that the MHR of PnPf2 is the important driver of protein-protein interactions. Nevertheless, our proposed model indicates that PnPf2 may form a homodimer *in situ* (**Figure 2A**), however these interactions may be occluded by the presence of the *Zn*-finger domain.

**Figure 2:**
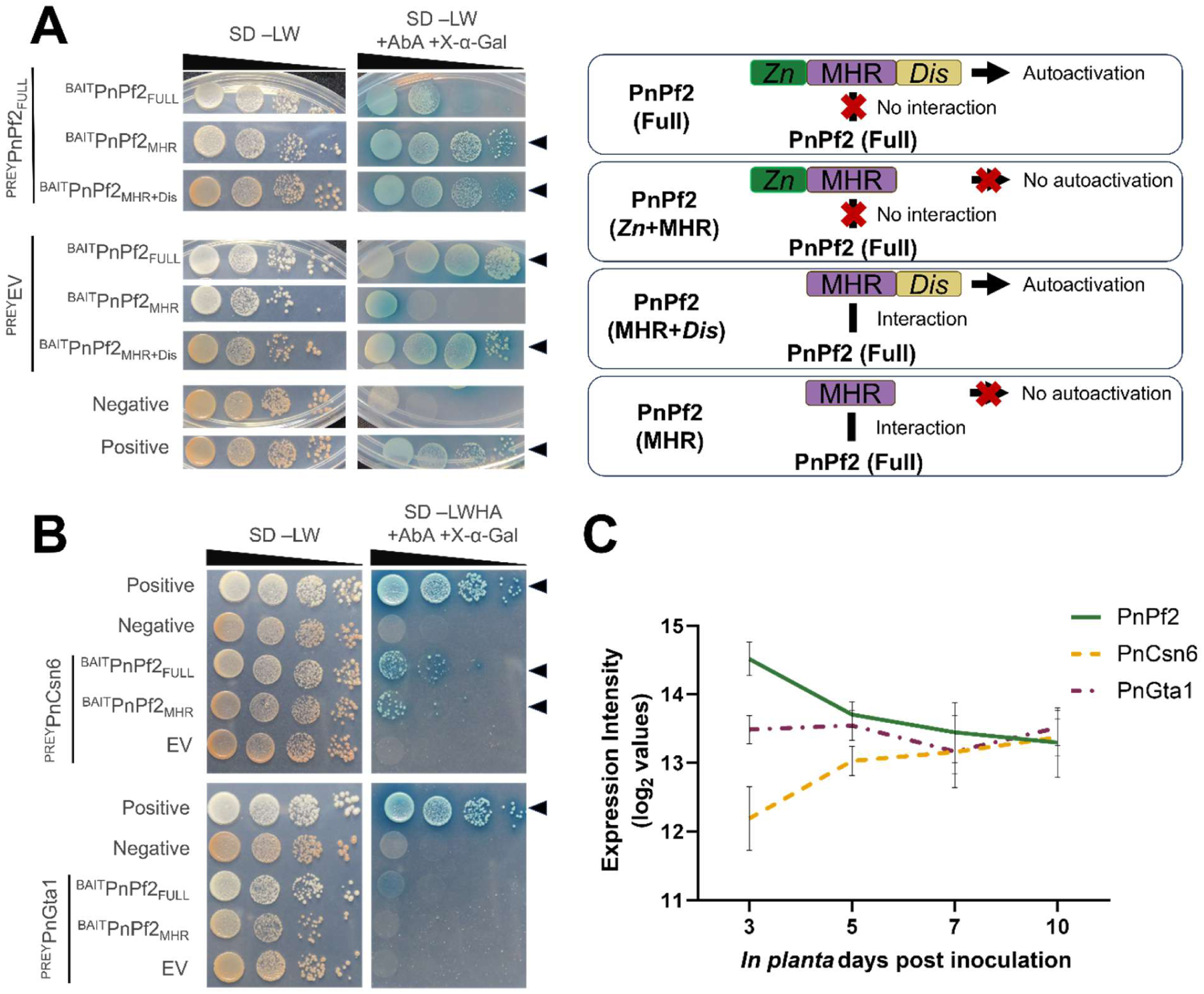
Yeast-2-hybrid (Y2H) assays reveal the PnPf2 MHR as the driver of protein-protein interactions. *S. cerevisiae* strains Y2HGold and Y187 carrying bait and prey vectors, respectively, were mated and grown on non-selective and selective media with serial-gradient cell concentrations. **(A)** PnPf2 homodimerises in a Y2H assay. Bait PnPf2 strains carry the GAL4 DNA binding domain translationally fused to various truncations of PnPf2: full-length PnPf2 (^BAIT^PnPf2_FULL_), MHR-only (^BAIT^PnPf2_MHR_) and MHR+*Dis* (^BAIT^PnPf2_MHR+Dis_), and were tested against the GAL4 activation domain translationally fused to full-length PnPf2 prey. The pGADT7 empty vector (EV) only (^PREY^EV) was used to test for autoactivation of PnPf2 against *only* the GAL4 activation domain. The full range of PnPf2-truncation Y2H tests are available in **Supplemental Figure S3**. Our proposed model of Y2H PnPf2 occlusion shown in the right panel. **(B)** PnPf2 interacts with full-length PnCsn6 in targeted Y2H but not with full-length PnGta1. GAL4 activation domain was translationally fused to full-length PnCsn6 (^PREY^Csn6) or PnGta1 (^PREY^Gta1) and tested against GAL4 DNA binding domain translational fusions of full-length (^BAIT^PnPf2_FULL_) and MHR-only truncated PnPf2 (^BAIT^PnPf2_MHR_), as well as the pGADT7 empty vector (EV). For both panels, growth on the selective media is indicative of interaction between the tested product and is highlighted with black arrows. Y2H assay negative and positive controls, from the Matchmaker Gold Y2H system, are also shown within each panel. **(C)** *In planta* microarray gene expression profile of *PnCsn6* and *PnGta1*, compared to *PnPf2*, over a 10-day time course (Ipcho et al., 2012).

### PnPf2 MHR drives protein-protein interactions with putatively co-localising nuclear proteins in yeast-2-hybrid

Previous yeast-1-hybrid experiments indicated PnPf2 may require co-regulators for efficient DNA binding (John et al., 2024). To uncover putative PnPf2 binding partners, we employed a similar Y2H approach as our PnPf2-homodimerisation assays, this time utilising the MHR PnPf2 as bait against a cDNA GAL4-AD fusion protein library of genes from SN15. Screening of our generated Y2H cDNA GAL4-AD fusion prey library against the PnPf2 MHR bait resulted in 380 positive blue colonies on the stringent selective X-α-Gal-containing media. To narrow down the list of putative interacting partners, we developed a high throughput sequence-screening method (see experimental procedures) which resulted to the identification of 64 unique positive interactions with PnPf2 MHR (^BAIT^PnPf2_MHR_) (**Supplemental Table S3**).

As PnPf2 has been shown to be localised within the nucleus (John et al., 2024), we shortlisted our candidate list to focus only on likely interactors with PnPf2 required for its DNA-binding/gene-regulatory roles. Therefore, the 64 potential interacting partners identified with our library-scale Y2H were shortlisted to six nuclear-localising proteins. Of these six candidates, two had characterised orthologues in other plant pathogenic fungi and were implicated in virulence (Lu et al., 2014, Chen et al., 2023, Lin et al., 2024): the COP-9 signalosome-associated subunit protein PnCsn6, and the putative Zn_2_Cys_6_-containing TF PnGta1. To validate interaction of these two PnPf2 MHR-binding candidates, we re-cloned each of the two protein-coding sequences of full-length PnCsn6 and PnGta1 into the pGADT7 prey vector to create new variants ^PREY^PnCsn6 and ^PREY^PnGta1. We performed direct-targeted Y2H assays with ^PREY^PnCsn6 and ^PREY^PnGta1 against the PnPf2 baits; both with PnPf2 full-length (^BAIT^PnPf2_FULL_) and the dimerisation-domain containing MHR (^BAIT^PnPf2_MHR_). We observed growth of blue colonies for the Csn6-PnPf2 interactions against full-length PnPf2 ^BAIT^PnPf2_FULL_ and the MHR domain-only PnPf2 ^BAIT^PnPf2_MHR_, indicative of protein-protein interaction in the Y2H assay (**Figure 2B**). Despite Y2H assays only being semi-quantitative (Estojak et al., 1995), the reduced colony growth at lower titres suggested that the interaction of PnCsn6 and PnPf2 was weaker than that of the Y2H positive control interactions, relative to total colony count in the assay. We were unable to detect growth and, therefore, unable to confirm PnPf2 protein-protein interaction for PnGta1 using the targeted Y2H assay (**Figure 2B**).

Since the Pf2 orthogroup is conserved in Ascomycetes (John et al., 2022), the phylogenies of Gta1 and Csn6 orthologues were analysed. Previous transcriptomic data indicate both *PnCsn6* and *PnGta1* are highly expressed in SN15 during *in planta* infection (Ipcho et al., 2012) but lack the classical “high to low” expression pattern observed for PnPf2, and other TFs implicated in necrotroph virulence (John et al., 2021, Verdonk et al., 2025) (**Figure 2C**). Both Csn6 and Gta1 orthologues were identified in all analysed fungal species, except for the absence of a Gta1 orthologue in publicly available genomes of the wheat pathogen *Zymoseptoria tritici*. Csn6 orthologues had a conserved functional COP9-signalosome domain that covered most of our identified protein sequences (**Figure 3A**). The conserved fungal COP9 signalosome assembles in a complex manner within fungi (Bakti et al., 2023), and is implicated in fungal development and secondary metabolite production (Maytal-Kivity et al., 2003, He et al., 2005, Mundt et al., 1999, Busch et al., 2003), so a Csn6 role with the carbon-metabolism regulator PnPf2 is conceivable.

**Figure 3:**
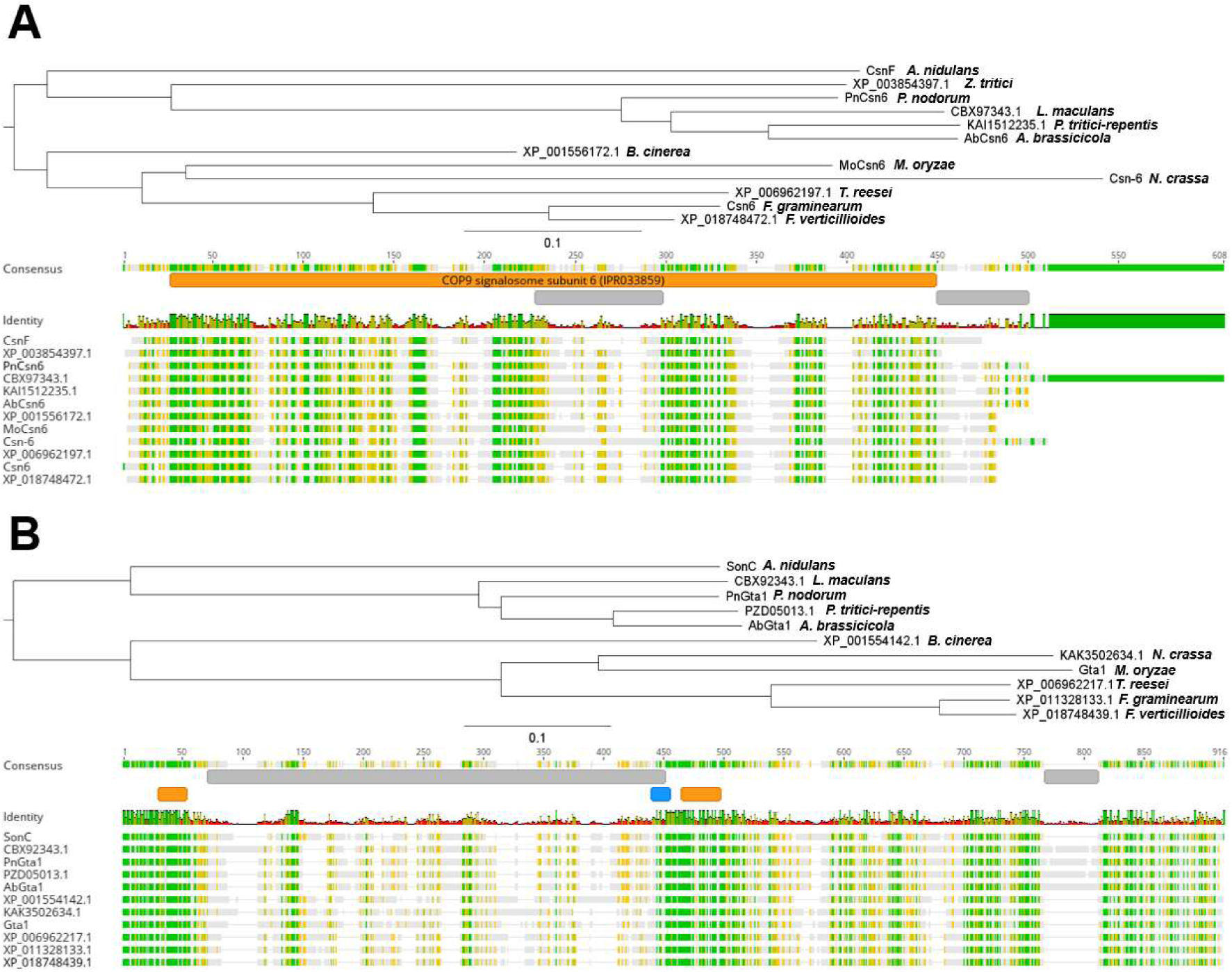
Phylogenetic analysis of (**A**) Csn6 and (**B**) Gta1 protein orthologues in 11 – 12 fungal species. Neighbour-Joining phylogenetic tree and sequence alignment of each orthologue are shown. The scale bars under the phylogenetic trees indicate substitutions per site. Orange bars indicate conserved putative functional domains, and the InterPro ID is in brackets for the Csn6 orthologue. Grey bars indicate disordered domains. Blue is the nuclear localisation signal in Gta1 orthologues. The domain boundaries in the aa sequence alignment are relative to the *P. nodorum* orthologues. GenBank accessions are provided as names for uncharacterised orthologues.

Despite being unable to validate the PnGta1-PnPf2 interaction, the protein architecture of the Gta1 orthologues is interesting. Gta1 orthologues feature a highly conserved N-terminal domain followed by a large, poorly conserved disordered region, then a unique conserved Zn_2_Cys_6_ domain first highlighted in the characterised Gta1 orthologue SonC from the model filamentous fungus *Aspergillus nidulans* (Larson et al., 2014) (**Figure 3B**). The N-terminus of Gta1 orthologues featured a conserved region resembling a C_2_HC zinc finger domain with many positively charged residues likely to promote nucleic acid binding (Cherstvy, 2009). However, just as the predicted Zn_2_Cys_6_ domain in SonC, the distance between the core cysteine/histidine residues in the putative C_2_HC domain (CX_2_CX_14_HX_4_C) is larger than typically found in characterised C_2_HC zinc finger proteins (CX_2_CX_12-14_HX_3_C) (Akhtar and Becker, 2001, Bijlmakers et al., 2016) (**Supplemental Figure S4A**). Despite the differences in motif from validated zinc finger domains, the zinc ion-coordinating residues in PnGta1 were confidently projected to be in the two putative zinc finger domains in structural predictions (**Supplemental Figure S4B**).

### PnCsn6 is required for vegetative growth, asexual reproduction and virulence in *P. nodorum*

As our targeted Y2H indicated that PnCsn6 directly interacts with PnPf2, we further investigated the role of *PnCsn6* in *P. nodorum* through phenotypic characterisation of a *PnCsn6*-deletion mutant. Concurrently, given its implication in fungal pathogenicity (Lu et al., 2014), we also investigated the potential role of PnGta1 in *P. nodorum* virulence. We generated two independent single-copy, deletion mutants of *PnCsn6* and *PnGta1* in *P. nodorum* SN15, along with accompanying ectopic marker-carrying mutants. Compared to SN15, deletion of *PnCsn6* in the mutants *pncsn6-3* and *pncsn6-5* resulted in reduced vegetative colony size and completely abolished pycnidiospore production on nutrient-rich V8PDA media (**Figure 4A**). In comparison, *PnGta1-*deletion mutants *pngta1-1* and *pngta1-14* had no discernible differences in vegetative growth phenotypes based on our assays. Similarly, *PnCsn6* is indispensable for *P. nodorum* pathogenicity, when tested in DLAs on the highly susceptible wheat cv. Halberd (**Figure 4B**). As we were unable to detect disease from the *PnCsn6* mutants using plug DLAs, we opted to significantly increase inoculum load by painting whole mycelia on the leaves. Additionally, we wondered if the reduced disease symptoms of the *pncsn6-*deletion mutants were due to the inability of the fungus to penetrate the host leaf epidermal layer. We then also pre-wounded the leaves prior to inoculation to assist in leaf-penetration. We noticed an increase in disease severity for the *pncsn6* mutants when painted mycelia is applied to both intact (unwounded) wheat leaves in DLAs, as well as for the pre-wounded wheat leaves – although each *pncsn6* mutant still showed reduced symptoms when compared to SN15 (**Figure 5A; Supplemental Figure S5**). Finally, the *pngta1* mutants showed no phenotypic difference from SN15 during infection based on our DLA observations. Taken together, these data indicate that PnCsn6 has a strong involvement in vegetative growth, asexual reproduction and is essential for *P. nodorum* virulence when infecting wheat, while the putative multi-zinc finger TF PnGta1 is dispensable for *P. nodorum* vegetative morphology and virulence.

**Figure 4:**
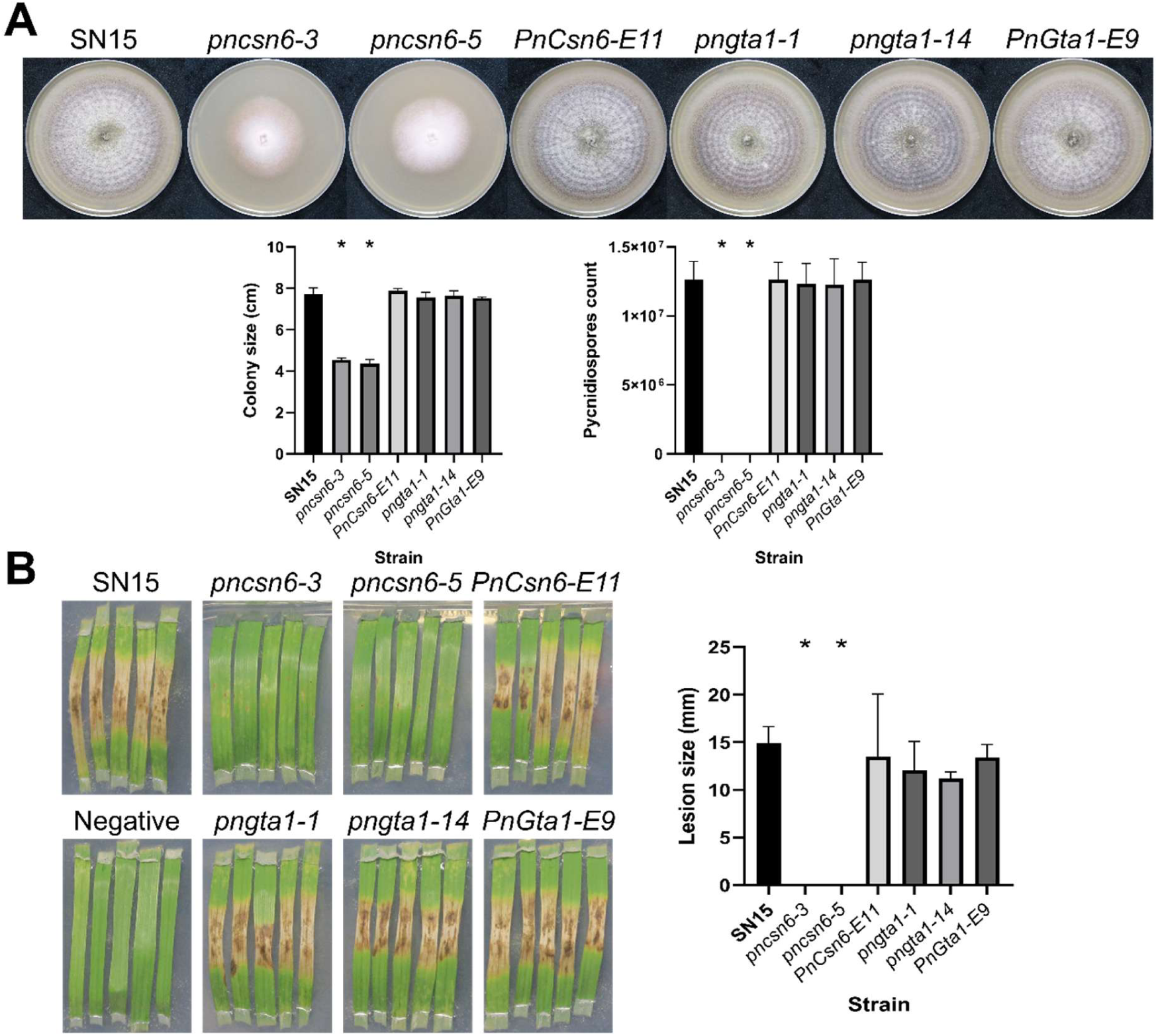
Characterisation of *PnCsn6*- and *PnGta1*-deleted mutants. (**A**) Vegetative growth of gene-deleted mutants showing clear growth deformities of *PnCsn6*-deleted mutants grown on V8PDA medium. The left bar graph shows colony sizes of each strain. The right bar graph shows the number of pycnidiospores produced by the strains per V8PDA plate. Ectopic mutants of *PnCsn6* and *PnGta1* (*PnCsn6-E11* and *PnGta1-E9*) showed SN15-like phenotypes. (**B**) Plug detached leaf assays (cv. Halberd) of *P. nodorum PnCsn6* and *PnGta1* mutant strains. No detectable lesions for both *pncsn6-3* and *pncsn6-5* indicate that PnCsn6 is required for virulence on wheat leaves. The right bar graph represents the average lesion size of each strain.

**Figure 5:**
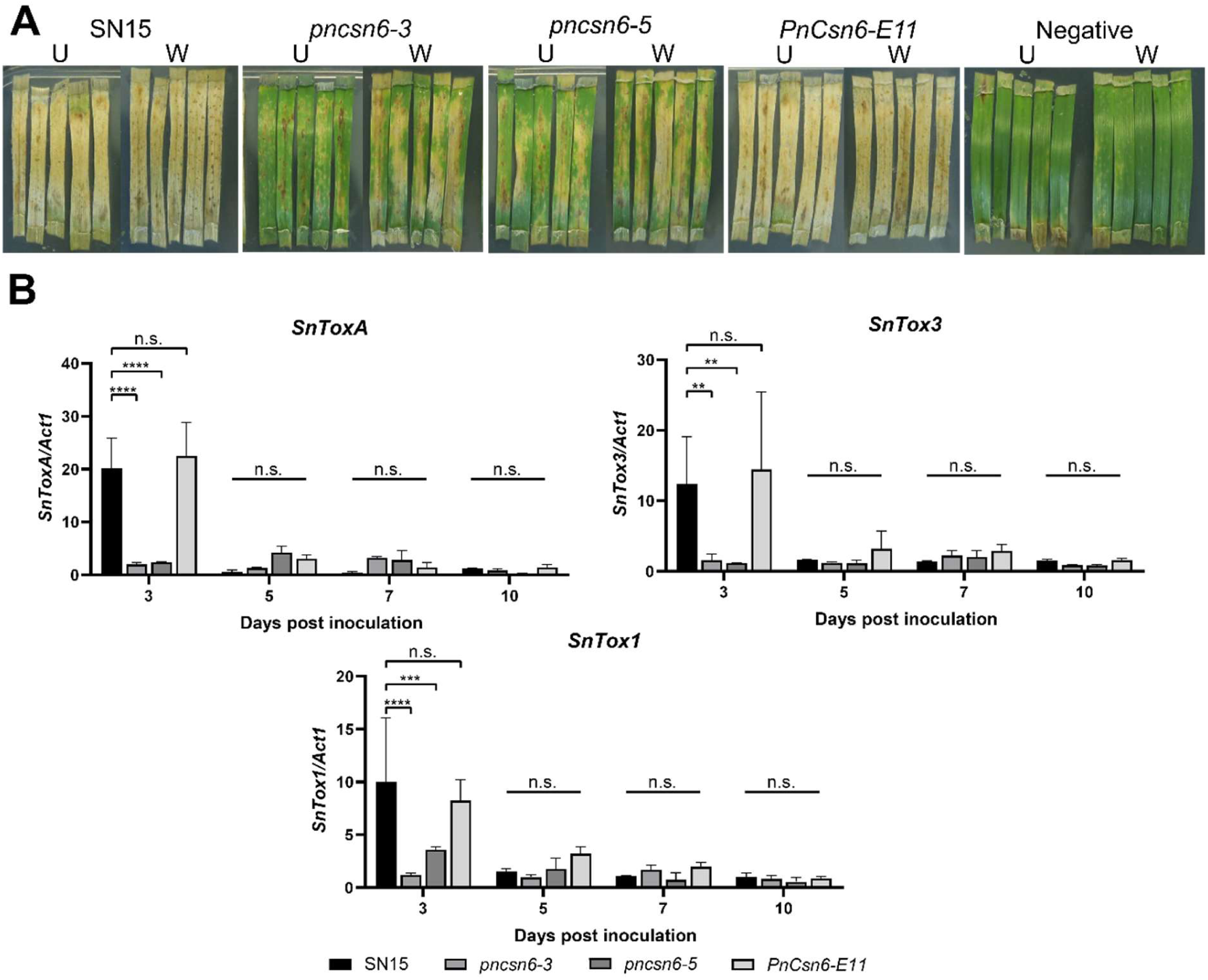
Impact of deletion of *PnCsn6* on wheat infection and NE gene expression. (**A**) Mycelia-painted DLAs (cv. Halberd) for *pncsn6-3* and *pncsn6-5*, along with ectopic *PnCsn6-E11*. Wounded leaves (W) show an improvement in disease progression over unwounded leaves (U) for *Csn6*-mutants, although overall reduced compared with SN15. Full time-course DLAs are available in **Supplemental Figure S5. (B)** Deletion of *PnCsn6* resulted in dysregulation of necrotrophic effector (NE) gene expression. Quantitative PCR of NE genes (*SnToxA, SnTox1* and *SnTox3*) normalised to the housekeeping actin (*Act1*) during infection of wheat (cv. Halberd) over three, five, seven and 10-days post inoculation. Adjusted *P-*values: ** ≤ 0.01, *** ≤ 0.001, **** ≤ 0.0001.

Given that both *PnCsn6*-deficient mutants exhibited reduced disease symptoms on wheat leaves, we investigated whether PnCsn6 has a direct role in regulating the expression of NEs in *P. nodorum*. To evaluate NE gene expression in *pncsn6-3* and *pncsn6-5*, mycelium-infected wheat leaves (cv. Halberd) were harvested at defined time points, and qRT-PCR was performed. At 3 days post inoculation, expression of *SnToxA* and *SnTox3* was significantly reduced - approximately 10-fold and 8-fold, respectively, compared with SN15 and the *PnCsn6-E11* ectopic strain (**Figure 4B**). Interestingly, we also observed that PnCsn6 has a direct effect on *SnTox1* expression, independent of PnPf2, as *PnCsn6-*mutants reduced detected *SnTox1* expression to ∼30% compared to SN15. We observed no other significant differences in NE gene expression for the later timepoints 5-, 7- and 10-days post inoculation. These results suggest that PnCsn6 has a direct role in NE gene regulation, perhaps through some pleiotropic effect during early infection of wheat within *P. nodorum*.

## DISCUSSION

Orthologues of the Pf2 transcription factor have been characterised in numerous phytopathogenic fungi and are established as requisite components for virulence (John et al., 2021). However, PnPf2 from *P. nodorum* is presently the sole Pf2 orthologue demonstrated to directly regulate gene expression through sequence-specific DNA binding (John et al., 2024). Our work here builds on this by demonstrating specific regulatory functions for individual domains within PnPf2, as well as unveiling the novel interacting partner PnCsn6, which shares virulence and effector-regulatory roles.

The PnPf2 MHR is highly conserved across fungal TFs (Shelest, 2008, Mayer et al., 2023), including the broadly conserved virulence-regulating Pf2 orthologues. Our results interrogating variable truncations of the PnPf2 gene highlighted the importance of the MHR domain, as truncation mutants lacking the MHR and putative 9aaTAD abolished detectable disease symptoms when infecting wheat and ablated necrotrophic effector (NE) gene expression as determined by qRT-PCR. Given the high level of sequence conservation in the MHR across Pf2 orthologues, it is likely that similarly important functions are retained (John et al., 2022). Predicted structural models of the MHR domain of PnPf2 superficially match the only experimental structures of MHR-containing proteins Cep3 (CBF3 subunit B): centromere/kinetochore-associated proteins within *Saccharomyces* and *Kluyveromyces* spp. (Dendooven et al., 2023, Lee et al., 2019, Yan et al., 2018, Zhang et al., 2018). Although aa sequence alignments of these proteins to PnPf2 are very low (≤10% aa identity), they share common structural motifs within the predicted MHR domain of PnPf2 and converge on α-Carbon root mean square deviation <5 Å, indicating distant homology (**Supplemental Figure S6**). Cep3 from *S. cerevisiae* contains a similar architecture to the TFs PnPf2 and Gal4 – consisting of an N-terminal zinc-finger DBD domain, followed by the MHR domain (Purvis and Singleton, 2008, Bellizzi et al., 2007). Cep3 is the major component of centromere binding factor 3 (CBF3), which assembles to targeted centromere DNA, often at the canonical epigenetic H3 histone markers, to enable chromosome segregation during cellular division (Guan et al., 2021, Mellone and Allshire, 2003, Choo, 2001). Indeed, Cep3 targets classical Zn_2_Cys_6_ TF major-groove DNA binding sites rich in CGG/CCG motifs (Schjerling and Holmberg, 1996, Shelest, 2017) much like PnPf2, as demonstrated with our DNA-motif enrichment searches using RNA-Seq (Jones et al., 2019) and ChIP-Seq (John et al., 2024). Such conservation of the MHR across global cell-division regulators may implicate Pf2 function beyond simple TF behaviour and epigenetic regulation during global cellular differentiation during host infection, although no such experimental evidence has been conducted to prove such a theory yet.

Intriguingly, we observed a direct reduction of *SnToxA* and *SnTox3* expression in the PnPf2-truncation mutants lacking the PnPf2 disordered region only (**Figure 1D**), and this disordered region is poorly conserved across pathogenic fungi (John et al., 2022, John et al., 2024). However, we do note that the specific portion of the PnPf2 disordered region required for NE gene expression (amino acids 497–529, surrounding the 9aaTAD) exhibits marginal conservation only among closely related fungal species. Moreover, intra-species polymorphisms within PnPf2 were most prevalent in this C-terminal disordered region. In combination, the disordered region of Pf2 may be important for regulating species- and strain-specific virulence factor genes, such as NEs *SnToxA* and *SnTox3* (McDonald et al., 2013), which is indeed the case for *P. nodorum*. The species-specific function mediated by the disordered region of the Pf2 orthologues may explain the low conservation of the region between species.

The requirement of homodimerisation for PnPf2 is consistent with other Zn_2_Cys_6_ domain-containing proteins, such as Cep3 and Gal4 from *S. cerevisiae* (Bellizzi et al., 2007, Hong et al., 2008), although the Zn_2_Cys_6_ domain of PnPf2 seemingly occluded interaction in our Y2H assay (**Figure 2A**). As the Y2H assay is based on fusion proteins of Gal4, the Zn_2_Cys_6_ TF PnPf2 translational fusions in Y2H may have unintended effects. Structural analysis of the MHR domain within Cep3 demonstrates a large dimerisation interface enabling homo- and heterodimerisation when assembling on DNA targets (Purvis and Singleton, 2008), whereby DNA-binding occurs through the centre of the protein complex, enabling the zinc-finger to target the dyad site and the MHR-interacting complex to externally face outward. Such an arrangement may explain why the translational fusion of PnPf2-Gal4-AD was unable to interact with any *Zn*-containing PnPf2-Gal4-DBD fusions, essentially a dual zinc-finger fusion, due to unintentional *Zn* DNA-interactions (**Supplemental Figure S2**). However, no occlusion was seen for the MHR-only Gal4-DBD fusion, enabling us to utilise this region for PnPf2 co-factor targets. Furthermore, we observed lethality in *S. cerevisiae* strain Y187 carrying ^PREY^PnPf2_FULL_, which limited our prey/bait combinations available for the Y2H assay. Indeed, this lethality for Y187 carrying full-length *PnPf2* may explain the inability to detect PnPf2 itself in the Y2H library screen using the MHR-only ^BAIT^PnPf2_MHR_. Lethality within the *S. cerevisiae* system may limit the detection of true interactions, a known limitation of the Y2H assay (Koegl and Uetz, 2007). To alleviate inherent Y2H issues, we did attempt to purify PnPf2 using heterologous expression in *E. coli*, either as full-length or truncation variants, however we were unable to reliably obtain purified protein for *in-vitro* experimentation presumably due to the intrinsic properties of the protein, as previously noted (John et al., 2024). Additionally, we were unable to reliably pull-down sufficient quantities of PnPf2 during our co-immunoprecipitation experiments for mass spectrometry identification of binding co-factors. Therefore, the use of alternative *in-vivo* methods, such as TurboID-mediated proximity labelling, should be considered for future validation and discovery of PnPf2-interacting partners, and such methods have been performed in other filamentous fungi (Hollstein et al., 2025).

Despite the inherent challenges of studying with PnPf2 in yeast systems, as we have previously noted (Jones et al., 2019), our results here have identified several biologically relevant, putative interacting partners of PnPf2. However, only PnCsn6 was validated as a PnPf2 binding partner using a two-way targeted Y2H. Csn6 is a subunit of the constitutive photomorphogenesis 9 (COP9) signalosome, a global signalling complex involved in development and stress tolerance (Braus et al., 2010, Dubiel et al., 2020), and its subunits are indispensable for virulence in other pathogens like *Fusarium graminearum* and *Magnaporthe oryzae* (Chen et al., 2023, Lin et al., 2024). PnCsn6 in *P. nodorum* appears to share similar functions to other Csn6-characterised orthologues. Interestingly, we previously characterised PnVeA from *P. nodorum* (Morikawa et al., 2024), and there are indications of interplay between the COP9 signalosome and the fungal-specific Velvet complex (Meister et al., 2019). Namely, the COP9 signalosome interacts with the conserved deubiquitinase UspA and down-regulates *UspA*, which in turn suppresses the accumulation of VeA in the fungus *Aspergillus nidulans* (Meister et al., 2019). PnPf2 may similarly interact with components of the COP9 signalosome, and potentially the velvet pathway, to coordinate shared cellular functions such as nutrient assimilation, carbon metabolism and fungal pathogenicity. This hypothesis needs to be validated in future studies.

The putative C_2_HC/Zn_2_Cys_6_ TF PnGta1 was identified as a PnPf2-interacting partner in a Y2H cDNA Gal4-AD fusion prey library screen. However, we were unable to validate this interaction in a targeted Y2H between PnPf2 and PnGta1, possibly due to the zinc finger domains in PnGta1 occluding the Y2H interaction, as we similarly suggested occurs with PnPf2 homodimerisation (**Figure 2A**). The gene knockout mutants of *PnGta1* were phenotypically indistinguishable from SN15 within our assays. In *M. oryzae*, Gta1 was required for virulence on rice, but the *Gta1* knockout mutant showed only minor growth defects (Lu et al., 2014). In *A. nidulans*, the Gta1 orthologue, SonC, was essential for fungal growth (Larson et al., 2014). As such, there appears to be a major functional divergence between Gta1 orthologues in different fungal species. Despite the lack of contribution of PnGta1 to the virulence of *P. nodorum*, Gta1/SonC orthologues feature unique conserved domains that may be of interest in future protein structural analyses.

Our PnPf2-bait Y2H library screen also identified conserved transcriptional initiation factors (**Supplemental Table S2**), including Taf13 (a subunit of TFIID) (Gupta et al., 2017) and Ess1 (a prolyl isomerase) (Hanes, 2014), indicating that PnPf2 may recruit essential components of the RNA polymerase II machinery. Additionally, interactions were detected with putatively nuclear-localising proteins PnMyb2 and PnRrm1. Although orthologues of these proteins are not well-characterised, related proteins containing Myb or RNA recognition motifs have been implicated in virulence and stress tolerance in other fungi (Kim et al., 2014, Zhang et al., 2020). This suggests PnMyb2 and PnRrm1 may have novel functions related to virulence in *P. nodorum,* although recent characterisation of similar domain-containing proteins did not show any effect on virulence (Verdonk et al., 2025). We were unable to validate the interactions of PnPf2 with PnTaf13, PnEss1, PnMyb2 or PnRrm1 in targeted Y2H assays (**Supplemental Figure S7**), suggesting our initial library-scale detected interactions may be transient or necessitate further validation through other methods.

Overall, our results here demonstrate that the PnPf2 MHR is critical to the regulation of fungal virulence, driving homodimerisation and direct recruitment of the transcriptional initiation machinery. Furthermore, we have functionally characterised the other domains of PnPf2 within the *P. nodorum* pathosystem, highlighting the role of a low-homology disordered region in controlling species-specific gene expression. The identification of a network of PnPf2-interacting partners provides a foundation for future investigations into the complex regulation of fungal pathogenesis. There is interest to identify the transcriptional regulators of PnPf2 itself, utilising DNA-investigation assays, such as yeast-1-hybrid (Reece-Hoyes and Marian Walhout, 2012), with fragments of the highly conserved 447 bp PnPf2 promoter as the bait, which could effectively identify these regulators and further unravel the transcriptional network controlling virulence and effector gene expression.

## EXPERIMENTAL PROCEDURES

### Strain and cultures

*P. nodorum* strains used in this study are outlined in **Table 2** and maintained on V8 potato dextrose agar (V8PDA) (150 ml L^−1^ Campbell’s V8 Juice, 3 g L^−1^ CaCO_3_, 10 g L^−1^ Difco PDA and 10 g L^−1^ agar) as described previously (Solomon et al., 2004a). *S. cerevisia*e strains Y2HGold and Y187 (Takara Bio) were grown in Yeast PDA (YPDA) or synthetic defined (SD) medium with selection as described in the Matchmaker® Gold yeast Two-Hybrid system user manual (Takara Bio). Yeast and bacterial strains used in this study can be found in **Supplemental Table S3.** All primers used in this study are described in **Supplemental Table S4.**

### Wheat infection assays

Detached leaf assay (DLA) was performed similarly to Solomon et al. (Solomon et al., 2004b), whereby approximately 4 cm of the first wheat leaves (cv. Halberd; *Tsn1, Snn1, Snn3*) were embedded adaxially into benzimidazole agar (75 mg/L benzimidazole and 15 g/L agar). *P. nodorum* strains used in this study were inoculated onto wheat leaves as either a mycelia/hyphal mixture (to ensure enough fungal mass for reliable RNA extraction), or as an agar plug (for consistent disease progression quantification). For mycelia/hyphal-mixture DLAs, 50 mg/mL of hyphae in 0.02% (v/v) Tween 20 solution were homogenised using TissueLyser II (QIAGEN) with a 1-mm diameter steel bead at 30 Hz for 10 sec, then was spread using a paintbrush prewet with 0.02% (v/v) Tween 20 solution. For (pre-)wounded leaves, wheat leaves were pierced through with a hypodermic 25G needle (Interpath) immediately prior to mycelia inoculation. Alternatively, for plug DLAs, approximately 2 x 2 mm agar plug was excised from a *P. nodorum* V8PDA plate and inoculated hyphae side down onto wheat leaves prewet with 0.02% (v/v) Tween 20 solution. The infection was left to develop for 3, 5, 7 and 10 days unless otherwise indicated. Negative treatments in all assays were performed with 0.02% (v/v) Tween 20 solution only. Effector expression assays were performed via qPCR as previously described (Rybak et al., 2017, John et al., 2024).

### Generation of *P. nodorum* PnPf2 truncation mutants

The nucleotide sequence of *PnPf2* (*SNOG_00649*) was acquired from the genome sequence and annotation of the Australian *P. nodorum* reference isolate SN15 (Bertazzoni et al., 2021). Truncations of PnPf2 were made according to the three domains defined by John et al. (John et al., 2024): a full-length *PnPf2* with the selectable marker at the 3’ UTR as a control strain (PnPf2_1-653_), a PnPf2 truncation missing the low homology disordered region (PnPf2_1-497_), and a PnPf2 truncation only containing the DNA-binding domain (PnPf2_1-63_). The putative PnPf2 9aaTAD was predicted by the webtool developed by Martin Piskáček (available at: https://www.med.muni.cz/9aaTAD/) (Piskacek et al., 2007), and PnPf2 truncations either side of this region (PnPf2_1-529_ and PnPf2_1-539_) were made. Truncation constructs were assembled using Fusion PCR (Hilgarth and Lanigan, 2020). *P. nodorum* strain SN15 was transformed via a PEG-mediated transformation as described previously (Solomon et al., 2004a). A single-copy integration mutant, as determined by qRT-PCR (Solomon, 2008), for each PnPf2-truncation was retained for further characterisation.

### Yeast-2-hybrid assays

Matchmaker® Gold Yeast Two-Hybrid System (Takara Bio) was used for the Y2H assay following the manufacturer’s protocol with PnPf2-specific modifications, detailed further here. Prey and bait plasmids were first cloned into and propagated in *Escherichia coli* strain TOP10 (ThermoFisher), then extracted using the QIAprep Spin Miniprep Kit (Qiagen) following the manufacturer’s protocol. Prey or bait plasmids were transformed into *S. cerevisiae* as described in Yeastmaker™ Yeast Transformation System 2 User Manual (Takara Bio). *S. cerevisiae* harbouring pGBKT7 plasmids were selected on SD media without tryptophan (SD-W), and pGADT7-containing *S. cerevisiae* were selected on SD media without leucine (SD-L). *PnPf2* exons were individually amplified and fused using fusion PCR, before being cloned into the bait vector pGBKT7 using HiFi Assembly (New England Biolabs, Ipswich, MA, USA). Various domain combinations of PnPf2 were subcloned from pGBKT7-PnPf2 (^BAIT^PnPf2_FULL_) by PCR of the desired domains along with the vector backbone. The purified PCR product was phosphorylated using T4 Polynucleotide Kinase (New England Biolabs), then ligated using T4 Ligase (New England Biolabs) according to the manufacturer’s protocols. All plasmids generated in this study were validated by MinION sequencing (Oxford Nanopore).

*P. nodorum in-vitro* RNA was isolated from SN15 grown in Fries 3 liquid media (30 g/L sucrose, 5 g/L ammonium tartrate, 1 g/L NH_4_NO_3_, 1 g/L yeast extract, 260 mg/L KH_2_PO_4_, 130 mg/L K_2_HPO_4_, 50 mg/L MgSO_4_·7H_2_O and 2 mL/L trace stock (227 mg/L CuSO_4_·5H_2_O, 167 mg/L LiCl, 34 mg/L H_2_MoO_4_, 72 mg/L MnCl_2_.4H_2_O, 80 mg/L CoCl_2_·4H_2_O in water)) for three days at 22°C. The SN15 cDNA library was synthesised using Make Your Own “Mate & Plate™” Library Kit (Takara Bio) following the manufacturer’s protocol and assembled into pGADT7-Rec digested with the *Sma*I restriction enzyme (New England Biolabs). All Y2H experiments were performed, in biological triplicate, as described in Matchmaker® Gold Yeast Two-Hybrid System User Manual, with *S. cerevisiae* Y2HGold carrying pGBKT7-PnPf2_MHR_ (^BAIT^PnPf2_MHR_) or pGBKT7-PnCsn6 (^BAIT^PnCsn6) used as bait for library-scale Y2H.

Targeted Y2H experiments were performed with candidate PnPf2-interacting prey proteins, re-cloned into pGADT7, and bait PnPf2: both as full-length PnPf2 (^BAIT^PnPf2_FULL_) and MHR-PnPf2 variant ^BAIT^PnPf2_MHR_, as well as the pGBKT7 (empty vector). All diploid *S. cerevisiae* cells were diluted to an OD600 of 1.0, serially diluted 1:10 in 0.9% *w/v* NaCl, and 20 µL spot plated onto SD media lacking leucine, tryptophan, histidine and adenine and supplemented with 1000 ng/mL aureobasidin (MedChemExpress) and 40 µg/mL X-α-galactose (SD-LWHA +AbA +X-α-Gal).

### Identification of PnPf2-interacting partners in yeast-2-hybrid

Positive colonies from the Y2H cDNA prey library screening against ^BAIT^PnPf2_MHR_ or ^BAIT^PnCsn6 were grown in batches of 10 positive colonies in 1 mL SD-LT liquid media for 24 hr at 30 °C at 200 rpm. All batch cultures were then pooled and grown in 250 mL of SD-L liquid media for a further 24 hr. *S. cerevisiae* plasmids were extracted using the Zymoprep Yeast Plasmid Miniprep II Kit (Zymo Research) following the manufacturer’s protocol. The extracted yeast prey candidate plasmids were transformed into *E. coli* TransforMax™ EPI300 (LGC Biosearch Technologies) and selected on lysogeny broth (LB) agar medium containing 100 µg/mL of ampicillin to remove endogenous yeast plasmids (2-micron) and bait plasmids. All *E. coli* cells were pooled by flooding the LB agar with LB liquid media containing 100 µg/mL of ampicillin and grown for 4 hr at 37 °C at 200 rpm and plasmid extracted using the QIAprep Spin Miniprep Kit (Qiagen). The extracted prey candidate plasmids were prepared for sequencing using the Rapid Barcoding Kit 24 V14 (Oxford Nanopore Technologies) and sequenced using MinION Mk1b (Oxford Nanopore Technologies).

The sequences were base called, and sequencing adapters were trimmed with Dorado v1.0.2 (https://github.com/nanoporetech/dorado). Sequencing reads were filtered with Nanoq v0.10.0 (Steinig, 2022) to retain reads with a quality score of Q15 and above, and a sequence length above 200 bp. BBDuk (https://sourceforge.net/projects/bbmap/) was used to trim the sequencing reads 30 bp upstream and downstream from the *Sma*I cut site in pGADT7-Rec, allowing for up to three bp mismatches. Trimmed sequencing reads over 50 bp in length were retained and mapped to the sequence of pGADT7-Rec using BBMap (https://sourceforge.net/projects/bbmap/) to remove the reads belonging to the plasmid backbone. The remaining unmapped reads were mapped to the *P. nodorum* SN15 genome (Bertazzoni et al., 2021) using Minimap2 (Li, 2018) and Samtools (Li et al., 2009). The mapped reads were located and visually confirmed with IGV (Robinson et al., 2011).

Localisation of candidate PnPf2 MHR interacting proteins was predicted using DeepLoc 2.1 (Ødum et al., 2024). As PnPf2 localises to the nucleus (John et al., 2024), the candidate PnPf2 MHR-interacting proteins were shortlisted to contain proteins predicted to localise to the nucleus. The shortlisted candidate PnPf2 MHR interacting proteins were interrogated using PhiB-BLAST (Urban et al., 2025) to identify characterised orthologues related to virulence. Shortlisted candidates with known hits to virulence-related orthologues were further characterised.

### Phylogenetic and sequence analyses

PnPf2, upstream and downstream sequences from a geographically diverse collection of 161 *P. nodorum* isolates (Pereira et al., 2021) were analysed by producing *de novo* draft assemblies using SPAdes v.3.14.1 (Bankevich et al., 2012). Draft assemblies were inspected using QUAST (Gurevich et al., 2013). Blastn was then performed using the canonical *PnPf2* sequence as a query. Each assembly produced a single, high-scoring hit. The blast hit coordinates were then used to extract the matching sequencing as well as 2000 bp up- and downstream. Extracted sequences were aligned using MAFFT v7.520 (Katoh et al., 2019) with the *maxiterate 1000* and *adjust-direction* options. The resulting multiple sequence alignment was visually inspected and trimmed to identical lengths. Protein sequences of Pf2 orthologues were retrieved from John et al. (John et al., 2024). A BLASTP search (Camacho et al., 2009) was utilised to find PnGta1 and PnCsn6 orthologues in the fungal species analysed by John et al. (John et al., 2024) (**Supplemental Table S1**). For orthologues in *A. brassicicola*, a BLASTN search was performed in the *Saccharomyces* Genome Database (Wong et al., 2023) with default parameters, except that the species was restricted to *A. brassicicola*. The identified hits were aligned with *PnCsn6* (*SNOG_13785*) and *PnGta1* (*SNOG_14970*) to predict the *AbCsn6* and *AbGta1* coding sequences, respectively.

Protein sequences were aligned using the default Clustal Omega algorithm on Geneious Prime 2024.0.4 (https://www.geneious.com/). Phylogenetic trees were constructed using the Neighbour-Joining algorithm with default settings and no outgroup. Disordered regions of PnCsn6 and PnGta1 were predicted using ADOPT (Redl et al., 2023). The conserved domain of PnCsn6 was annotated using InterPro (Blum et al., 2025). The putative nuclear localisation signal of PnGta1 was determined using NLStradamus (Nguyen Ba et al., 2009) with a 0.5 cutoff and a 4-state HMM static setting.

AlphaFold 3 (Abramson et al., 2024) was used to predict the protein structure of PnPf2 and PnGta1. Three zinc ions were added to predict the potential ion coordination of the putative zinc finger domains in PnGta1, and two zinc ions were added for the PnPf2 models. The predicted protein models were visualised in Open-Source PyMOL version 3.1 (https://github.com/schrodinger/pymol-open-source) and coloured using pymol-color-alphafold (https://github.com/cbalbin-bio/pymol-color-alphafold). Foldseek was used to identify structurally similar proteins to the PnPf2 MHR (van Kempen et al., 2024).

## ACKNOWLEDGEMENTS

The authors thank Dr Huyen T. T. Phan and Dr Johannes W. Debler for their assistance during laboratory experiments.

## AUTHOR CONTRIBUTIONS

<u>Shota Morikawa</u>: Investigation, Conceptualisation, Formal analysis, Data Curation, Methodology, Validation, Writing - original draft. Writing - review & editing. <u>Carlia Herbst</u>: Investigation. <u>Evan John</u>: Supervision. <u>Daniel Croll:</u> Investigation. <u>Carl Mousley</u>: Supervision. <u>Bernadette Henares</u>: Supervision. <u>Kar-Chun Tan</u>: Conceptualisation, Project administration, Resources, Supervision, Funding acquisition, Writing - review & editing. <u>Callum Verdonk</u>: Investigation, Methodology, Validation, Supervision, Writing - original draft, Writing - review & editing. All authors contributed to the finalised manuscript prior to submission.

## DATA AVAILABILITY

The data that support the findings of this study are openly available in-text or in the supporting information.

