## Supplementary data for "A conserved fungal transcription factor domain drives protein-protein interactions and necrotrophic effector-mediated virulence in *Parastagonospora nodorum*"

**Supplemental Table S1**: Fungal species and strains used for the phylogenetic analyses presented in-text.

| Species | Strain | Pairwise identity [positive] % to PnPf2 |
|---|---|---|
| *Alternaria brassicicola* | ATCC 96836 | 74.5 [85.3] |
| *Aspergillus nidulans* | FGSC A4 | 21.4 [37.9] |
| *Botrytis cinerea* | B05.10 | 35.2 [50.8] |
| *Fusarium graminearum* | PH-1 | 24.5 [36.5] |
| *Fusarium verticillioides* | 7600 | 24.3 [36.6] |
| *Leptosphaeria maculans* | JN3 | 74.3 [85.3] |
| *Magnaporthe oryzae* | 70-15 | 29.3 [44.7] |
| *Neurospora crassa* | FGSC 2489 | 30.8 [47.2] |
| *Parastagonospora nodorum* | SN15 | 100 |
| *Pyrenophora tritici-repentis* | M4 | 71.5 [82.0] |
| *Trichoderma reesei* | QM6a | 29.9 [46.4] |
| *Zymoseptoria tritici* | IPO323 | 39.3 [54.5] |

**Supplemental Table S2**: Sixty-four putative PnPf2-interacting partners identified in Y2H cDNA library screen against PnPf2 middle homology region (sorted by locus number). Bolded rows indicate the candidates predicted to localise in the nucleus and tested in this study.

| Gene Identifier | Function/annotation |
|---|---|
| SNOG_00117 | 37S ribosomal protein S25 |
| SNOG_00270 | Lipoyl synthase |
| SNOG_00387 | Uncharacterised protein |
| SNOG_00848 | Peptidyloprolyl isomerase |
| SNOG_00990 | Heme haloperoxidase |
| SNOG_01278 | Glyceraldehyde-3-phosphate dehydrogenase GPD1 |
| SNOG_01936 | Large ribosomal subunit protein uL2 |
| SNOG_02221 | Cholestenol Delta-isomerase |
| SNOG_02498 | Large ribosomal subunit protein mL50 |
| **SNOG_02555** | **RNA Recognition Motif domain (PnRRM1)** |
| SNOG_03302 | RNA helicase |
| SNOG_03533 | Peptidylprolyl isomerase |
| SNOG_03650 | Ribosomal protein L38e |
| SNOG_03785 | 60S ribosomal protein L36 |
| SNOG_03888 | ATP phosphoribosyltransferase |
| SNOG_04100 | F-box domain-containing protein |
| SNOG_04135 | Proliferating cell nuclear antigen |
| SNOG_04428 | Inorganic pyrophosphatase |
| SNOG_04609 | Mitochondrial protein import protein MAS5 |
| SNOG_04647 | Protein kinase domain |
| SNOG_04649 | Bromo domain-containing protein |
| **SNOG_04857** | **Transcription initiation factor TFIID subunit 13 (PnTaf13)** |
| SNOG_04916 | PPM-type phosphatase domain |
| SNOG_04923 | Mitochondrial outer membrane protein porin |
| SNOG_05148 | Anaphase-promoting complex subunit 4 WD40 |
| SNOG_05719 | Small ribosomal subunit protein uS2 |
| SNOG_06036 | RNA-binding S4 domain |
| SNOG_07147 | TPR domain-containing protein |
| SNOG_07852 | Actin-like ATPase domain-containing protein |
| SNOG_07942 | Heterokaryon incompatibility domain |
| SNOG_08021 | Uncharacterised protein |
| SNOG_08046 | Bacterial surface antigen D15 |
| SNOG_08966 | NADH dehydrogenase ubiquinone iron-sulphur protein 4 |
| SNOG_09012 | Large ribosomal subunit protein uL23 |
| SNOG_09371 | NADH-ubiquinone oxidoreductase |
| SNOG_09416 | Glucose-6-phosphate 1-dehydrogenase |
| SNOG_10409 | Large ribosomal subunit protein uL18 |
| SNOG_11098 | Sulphite oxidase |
| SNOG_11193 | Nucleoside diphosphate kinase |
| SNOG_11675 | Peptidase M20 dimerisation domain-containing protein |

| | |
|---|---|
| SNOG_11881 | beta-glucosidase |
| SNOG_11918 | phenylalanine--tRNA ligase |
| SNOG_12064 | Small ribosomal subunit protein uS10 |
| SNOG_12075 | Putative membrane protein |
| SNOG_12085 | aldose 1-epimerase |
| SNOG_13370 | Uncharacterised protein |
| **SNOG_13398** | **Myb domain Transcription Factor (PnMyb2)** |
| SNOG_13582 | DUF2415-containing domain |
| **SNOG_13763** | **Peptidyl-prolyl cis-trans isomerase (PnEss1)** |
| **SNOG_13785** | **COP9 signalosome complex subunit 6 (PnCsn6)** |
| SNOG_14474 | Chitinase |
| SNOG_14739 | AP-3 complex subunit mu |
| **SNOG_14970** | **Zn2Cys6 domain Transcription Factor (PnGta1)** |
| SNOG_15174 | Yeast cell wall synthesis Kre9/Knh1-like N-terminal domain |
| SNOG_15454 | Major facilitator superfamily |
| SNOG_15478 | DUF1996-containing protein |
| SNOG_15488 | Malate dehydrogenase Mdh1 |
| SNOG_15657 | 40S ribosomal protein S8 |
| SNOG_15879 | 6-phosphogluconate dehydrogenase |
| SNOG_16121 | Heat shock protein |
| SNOG_16163 | Cysteine desulphurase |
| SNOG_16543 | T-complex protein 1 subunit delta chaperone |
| SNOG_42246 | Uncharacterised protein |
| SNOG_42379 | F-box domain-containing protein |

**Supplemental Table S3**: Strains used in this study.

| Strain | Description | Source |
|---|---|---|
| | <u>*S. cerevisiae* strains used</u> | |
| Y187 | Y2H strain. Further information found in Matchmaker Gold Yeast Two-Hybrid System User Manual. | Harper et al. 1993*. Takara Bio |
| Y2HGold | Y2H strain. Carries the *AUR1-C* ($AbA^R$) reporter. Further information found in Matchmaker Gold Yeast Two-Hybrid System User Manual. | Nguyen, Unpublished. Takara Bio |
| | <u>*E. coli* strains used</u> | |
| EPI300 | Used in this study for propagating plasmids. | Epicentre |
| Top10 | Used in this study for cloning and propagating plasmids. | ThermoFisher Scientific |

* Harper JW, Adami GR, Wei N, Keyomarsi K, Elledge SJ. The p21 Cdk-interacting protein Cip1 is a potent inhibitor of G1 cyclin-dependent kinases. Cell. 1993 Nov 19;75(4):805-16. doi: 10.1016/0092-8674(93)90499-g. PMID: 8242751.

**Supplemental Table S4**: Primers used in this study

| Name | Sequence | Purpose |
|---|---|---|
| PnPf2_Zn2_R | TACAAAATTCATCCCAATACTCATCATC AGAGCACCTTGGCCCTGCTCCC | PnPf2 truncations |
| PnPf2_Dis2_R | TACAAAATTCATCCCAATACTCATCATC AAGCAGGGAGTGTCTGGG | PnPf2 truncations |
| PnPf2_Full2_R | AAAATTCATCCCAATACTCATCATCATT GCTTGAACATCATAGATGGGTC | PnPf2 truncations |
| PnPf2_DisBTail_R | ATACAAAATTCATCCCAATACTCATCAT CACTGAGCAGCAAGCCGGC | PnPf2 truncations |
| PnPf2_DisCTail_R | ATACAAAATTCATCCCAATACTCATCAT CAAGACGGGTACGATGGCG | PnPf2 truncations |
| PnPf2_pTef_3F | CTTAACTTGGGCGGTGATTCTGCTGTC TCGCTCTTGTTGATGACGCCGC | PnPf2 truncations |
| tTef_SCx3_R | TGATGATGAGTATTGGGATGAATTTTG TATGCACGC | PnPf2 truncations |
| pGADT7_Pf2_F | CGCTCATATGGCCATGGAGGCCAGTGA ATTATGTCGTCCAGCAGTACCAC | Y2H PnPf2 |
| pGADT7_Pf2_R | ATCTGCAGCTCGAGCTCGATGGATCTC ATTGCTTGAACATCATAGATGGG | Y2H PnPf2 |
| pGBKT7_Pf2_F | GACCTGCATATGGCCATGGAGGCCGAA TTCATGTCGTCCAGCAGTACCAC | Y2H PnPf2 |
| pGBKT7_Pf2_R | GCGGCCGCTGCAGGTCGACGGATCCTC ATTGCTTGAACATCATAGATGGG | Y2H PnPf2 |
| pGBK_F_linear | ATAACTAGCATAACCCCTTG | Y2H bait construction |
| pGBK_R_linear | GGCCATATGCAGGTCCTCCT | Y2H bait construction |
| pGADT7-AD_linear_FWD | GGGTGGGCATCGATACGGGA | Y2H prey construction |
| pGADT7-AD_linear_REV | GTGGAATTCACTGGCCTCCA | Y2H prey construction |
| pGADT7_Screen_R | GAAAGAAATTGAGATGGTGCAC | Y2H prey screen |
| pGBKT7_Screen_R | TTCGCCCGGAATTAGCTTGG | Y2H bait screen |
| pG_Y2Hvec_screen_F | TAATACGACTCACTATAGGG | Y2H vector screen |
| PnPf2_DisY2H_F | TCAACGATCAGCATGTCAGG | Y2H PnPf2 truncations |
| PnPf2_MHRY2H_F | CTCCTTCCCAATGGACTGG | Y2H PnPf2 truncations |
| PnPf2_MHRY2H_R2 | TTAAGCAGGGAGTGTCTGGGG | Y2H PnPf2 truncations |
| PnPf2_ZnY2H_R2 | TTAACCCTGAGCGCGTGCGAA | Y2H PnPf2 truncations |
| PnCsn6_GAD_F | TGGAGGCCAGTGAATTCCACGAAGCTT CACATAATCCGCTG | Y2H PnCsn6 |
| PnCsn6_GAD_R | TCCCGTATCGATGCCCACCCTCACCAT GCAGCTCCACTAC | Y2H PnCsn6 |
| PnGta1_GAD_F | TGGAGGCCAGTGAATTCCACCGCAGAC CCAACCAATTGAT | Y2H PnGta1 |
| PnGta1_GAD_R | TCCCGTATCGATGCCCACCCTCAGTCAC CCCGGATAATTT | Y2H PnGta1 |
| PnMyb2_exon1_R | ATTTGTGTTGAGATGGAACGAC | Y2H PnMyb2 |
| PnMyb2_exon2_F | TTAGCAACACTTCTAGTAGCATC | Y2H PnMyb2 |
| PnMyb2_GAD_F | TGGAGGCCAGTGAATTCCACATGCTGGTCCTCAGCACGTCAA | Y2H PnMyb2 |
| PnMyb2_GAD_R | TCCCGTATCGATGCCCACCCCTACATGATGTTCTTGAGGTTCATC | Y2H PnMyb2 |
| Taf13_exon1_R | TAGAAGCCTTCCTCCTCGC | Y2H PnTaf13 |
| Taf13_exon2_F | AGTCTTGTGTGGGAAACTGC | Y2H PnMyb2 |
| Taf13_GAD_F | TGGAGGCCAGTGAATTCCACATGACAGAGGTGCGCATGC | Y2H PnMyb2 |
| Taf13_GAD_R | TCCCGTATCGATGCCCACCCTCATGCACTCATGACCGTGC | Y2H PnMyb2 |
| Ess1_exon1_R | AGATCGCCCCCTTTACGTG | Y2H PnEss1 |
| Ess1_exon2_F | TGGATTCTTTGGCCACGGAG | Y2H PnEss1 |
| Ess1_GAD_F | TGGAGGCCAGTGAATTCCACATGTCTTCAGAGGAGACAGGTC | Y2H PnEss1 |
| Ess1_GAD_R | TCCCGTATCGATGCCCACCCCTACCTCTGAATCAAATGCAG | Y2H PnEss1 |
| 02555_exon1_R | CTTTTGATTATTGAAAGTCGCAATG | Y2H PnRrm1 |

| | | |
|---|---|---|
| 02555_exon2_F | GCCGACGGCAGAGTTCTG | Y2H PnRrm1 |
| 02555_GAD_F | TGGAGGCCAGTGAATTCCACATGCCGGACTCGACCTCA | Y2H PnRrm1 |
| 02555_GAD_R | TCCCGTATCGATGCCCACCCCTAGTAACGCCTTCGTGGTC | Y2H PnRrm1 |
| PnCsn6_3F | CTTAACTTGGGCGGTGATTCTGCTGTCT CGTCAGTGTCGGCAGCAAAGG | PnCsn6 deletion |
| PnCsn6_3R | GTATTCAATTTCCGACTTTTGCG | PnCsn6 deletion |
| PnCsn6_3R2 | TGCGTCAGAGGTCTTTGGG | PnCsn6 deletion |
| PnCsn6_3Screen_R | CTGAGGGAAATGTGATTCGG | PnCsn6 deletion |
| PnCsn6_5F | CAGCAACTTACAAGTGAGCAG | PnCsn6 deletion |
| PnCsn6_5F2 | AGATCATATGCTAACCTGGC | PnCsn6 deletion |
| PnCsn6_5R | AGCGCGTGCATACAAAATTCATCCCAAT ACGATGATGGCACCAATCGAG | PnCsn6 deletion |
| PnCsn6_5Screen_F | AGATCTGCATGCAAGCCGG | PnCsn6 deletion |
| PnGta1_3F | AGCGCGTGCATACAAAATTCATCCCAATA CTCGGGCTTATCACGATTGAG | PnGta1 deletion |
| PnGta1_3R | CGAGTGCCTCCTCACAATTC | PnGta1 deletion |
| PnGta1_3R2 | CTCCTCACAATTCTAACACG | PnGta1 deletion |
| PnGta1_3Screen_R | CCAGCGACAGAAGAAGACG | PnGta1 deletion |
| PnGta1_5F | CTCCTCCATCAGTGTAACAG | PnGta1 deletion |
| PnGta1_5F2 | GTTTAGTCTCTGAAGCTGCG | PnGta1 deletion |
| PnGta1_5R | CTTAACTTGGGCGGTGATTCTGCTGTCTC GGCCCAAGGATTCAAGCAGC | PnGta1 deletion |
| PnGta1_5Screen_F | TGTCTAAGGGAAGAGAAGTC | PnGta1 deletion |
| ToxA_qPCR_F | CGATCCCGGTTACGAAATC | qPCR NEs |
| ToxA_qPCR_R | TTGACATGCAGCTTCCCTG | qPCR NEs |
| Tox1_qPCR_F | TGGTCTTGTCAGTAGCCTTTGC | qPCR NEs |
| Tox1_qPCR_R | TCCTGGAGTATGGCAAATTG | qPCR NEs |
| Tox3_qPCR_F | AATGTCGACCGTTTTGACC | qPCR NEs |
| Tox3_qPCR_R | GGTTGCCGCAGTTGATATAA | qPCR NEs |
| tTef_Screen_F | CCATCATGCCACCAAAAGCG | Mutant copy-number validation |
| pTef_Screen_R | TACGTAGCAAGATAGACCGC | Mutant copy-number validation |
| tTef_R | GTATTGGGATGAATTTTGTATGCACGC | Mutant copy-number validation |
| pTef_F | CGAGACAGCAGAATCACCGC | Mutant copy-number validation |
| Actin_qPCR_F | AGTCGAAGCGTGGTATCCT | qPCR relative marker *Act1* |
| Actin_qPCR_R | ACTTGGGGTTGATGGGAG | qPCR relative marker *Act1* |
| Phleo_qPCR_F | ACTTCATCGCAGCTTGACTAAC | Mutant copy-number validation |
| Phleo_qPCR_R | TGATGAACAGGGTCACGTC | Mutant copy-number validation |
| HygD F | CGAAGAATCTCGTGCTTTCA | Mutant copy-number validation |
| HygD R | TCTTTGTAGAAACCATCGGC | Mutant copy-number validation |

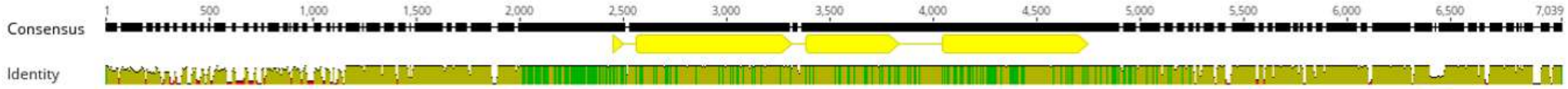


**Supplemental Figure S1**: Nucleotide sequence alignment of *PnPf2* and its surrounding genomic region in 161 isolates of *P. nodorum*. Yellow bars represent the coding sequence (exons) of PnPf2. Colours in “Identity” show sequence similarities in a 1-bp sliding window – Red: <30%; brown: 30 – <100%; green: 100%.

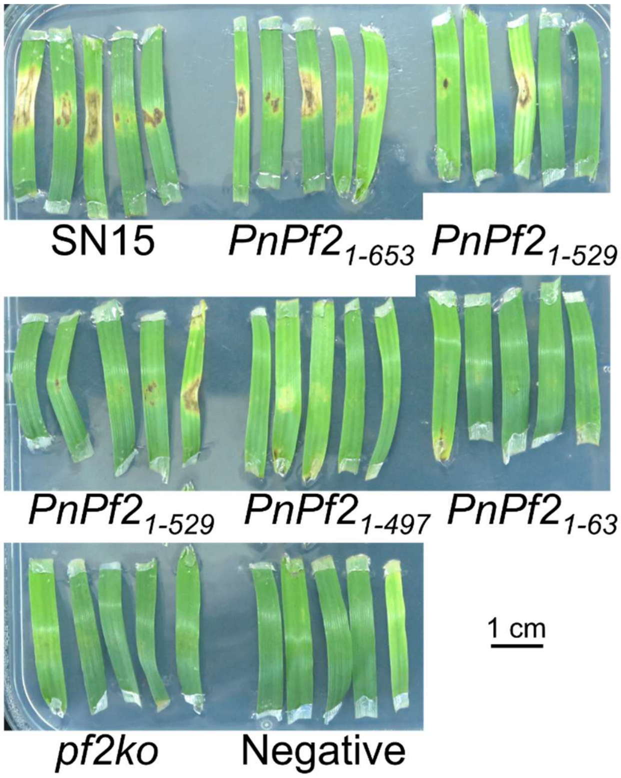


**Supplemental Figure S2**: Plug DLAs (cv. Wyalkatchem) of PnPf2-truncation carrying strains 10-dpi. SN15 and the *Dis*-deleted strain *PnPf2*$_{1\text{-}497}$ are also shown in-text in **Figure 1E**.

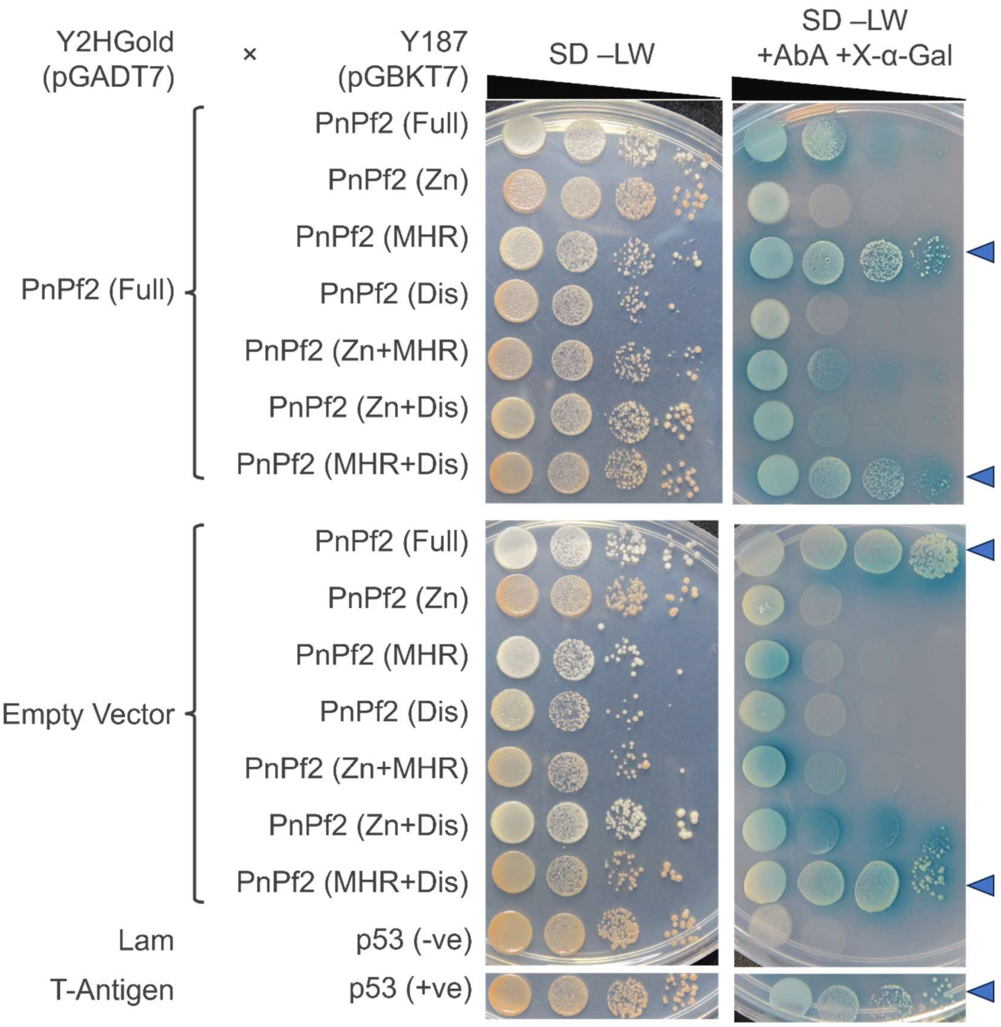


**Supplemental Figure S3**: Full Y2H results to complement in-text Figure 2. *S. cerevisiae* strains Y2HGold and Y187 carrying pGADT7 and pGBKT7 vectors, respectively, were mated and grown on non-selective (SD –LW) and selective (SD –LW +AbA +X-α-Gal) media with a gradient of cell concentrations. The top panel depicts GAL4 DNA binding domain in pGBKT7 was translationally fused to various truncations of PnPf2 and tested against GAL4 activation domain in pGADT7 translationally fused to PnPf2. The bottom row is the same assay, but the GAL4 activation domain was not fused to PnPf2 to test autoactivation. Growth on the selective media is indicative of interaction between the products of pGADT7 and pGBKT7 and is highlighted with blue arrows on the right. The bottom two rows are the positive (+ve) and negative (-ve) controls, respectively. Full – full-length protein; Zn – $Zn_2Cys_6$ DNA binding domain; MHR – middle homology domain; Dis – C-terminal disordered domain.

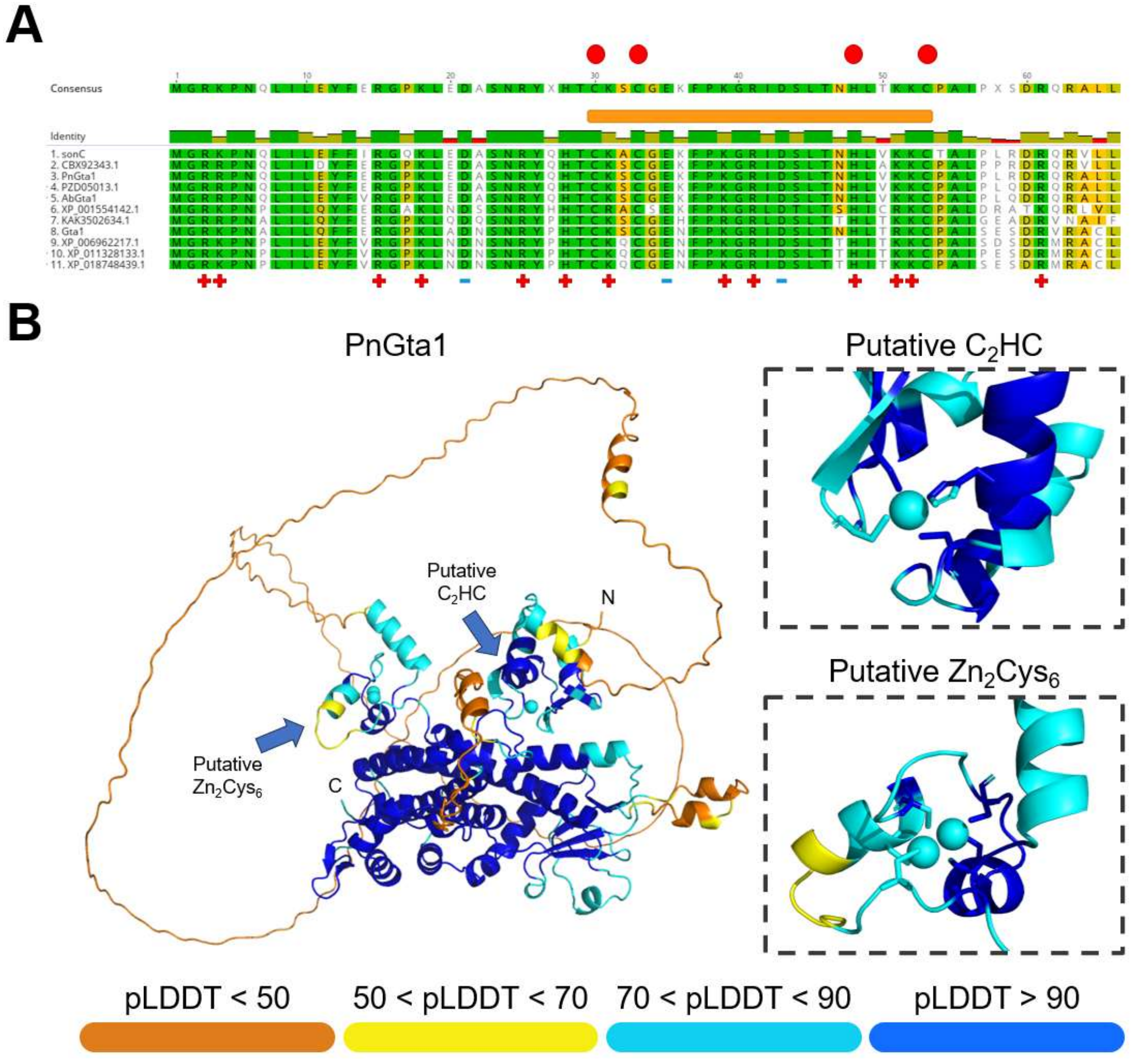


**Supplemental Figure S4**: Analysis of the putative zinc finger domains in Gta1 orthologs. (**A**) Amino acid sequence alignment of Gta1 orthologs (residues 1 – 66) showing the conserved N-terminus with the putative $C_2HC$ zinc finger. The red circles above the alignment show the conserved residues of the putative $C_2HC$ zinc finger domain. The orange bar represents the approximate boundaries of the domain. The symbols below the alignment show polarity of the amino acids: (+) positively charged; (-) negatively charged. (**B**) AlphaFold 3 protein structure prediction of PnGta1. The two panels on the right show a close-up of the two zinc finger domains and predicted zinc ion coordination, and the side chains of the coordinating residues are shown. The predicted PnGta1 model is coloured according to predicted local distance difference test (pLDDT) scores of the residues.

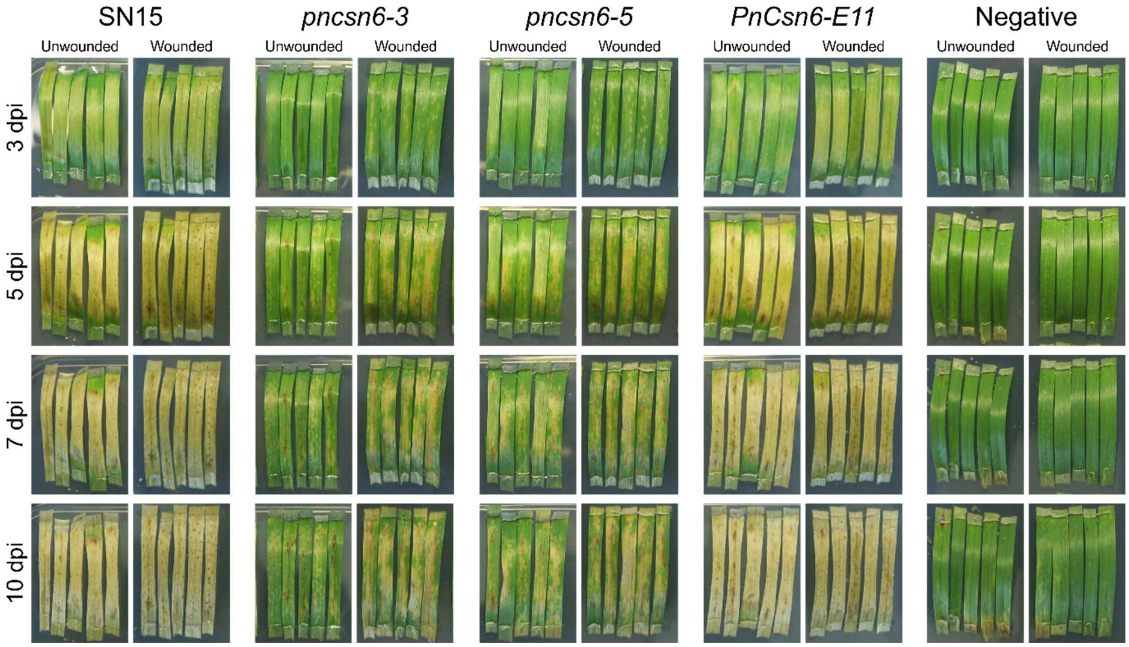


**Supplemental Figure S5:** Whole-mycelia (painted) inoculum DLAs (wheat cv. Halberd) for *PnCsn6* mutants at various timepoints. Wheat leaves were pre-wounded (right images) prior to application of mycelia. Significant reduction of disease symptoms are observed for *PnCsn6* mutants in DLA.

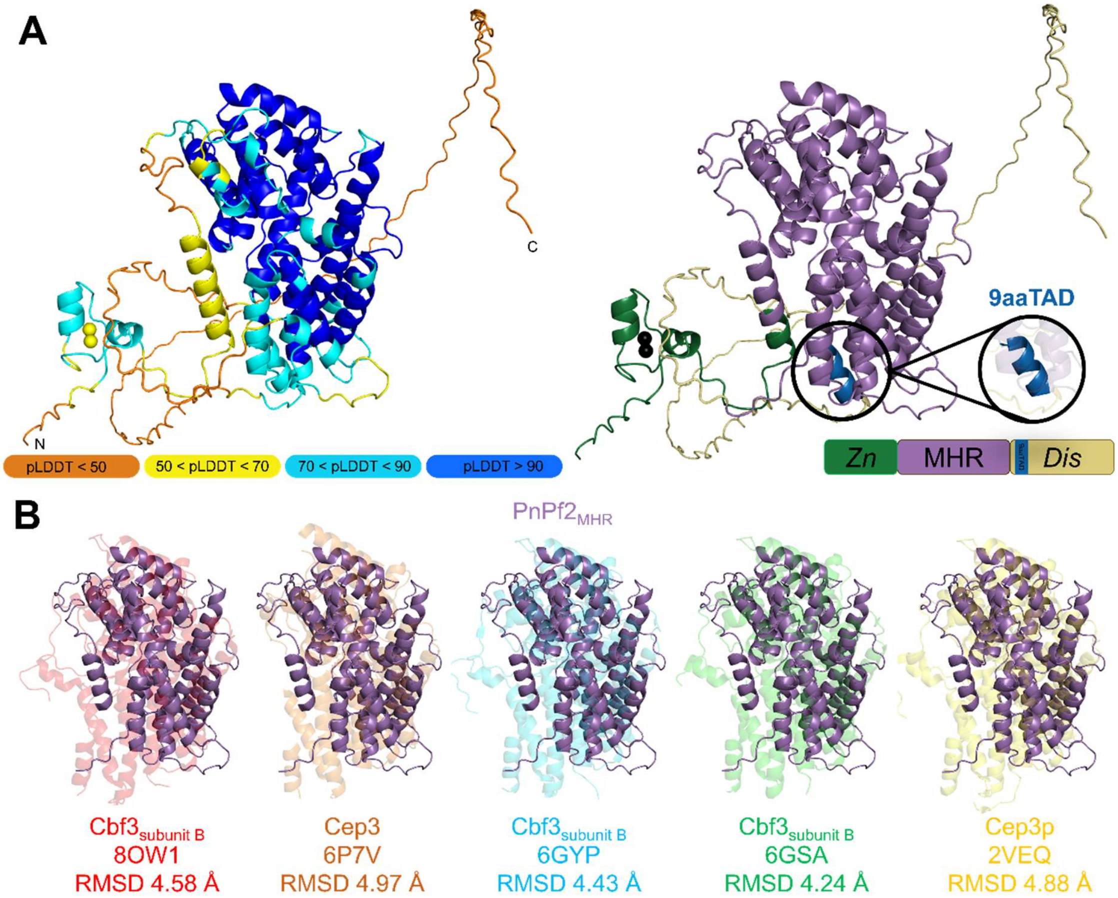


**Supplemental Figure S6:** Structural prediction models of PnPf2. **(A)** The PnPf2 Alphafold3 model model coloured by predicted local distance difference test (pLDDT) (Left), while the right panel shows the relative arrangement of the PnPf2 domains as described in-text: *Zn*, $Zn_2Cys_6$ domain (green); MHR, middle homology region (purple); *Dis*, C-terminal disordered domain (yellow). Predicted 9aaTAD coloured in blue. **(B)** Structural alignment of (experimentally resolved) zinc-finger and MHR-containing proteins from *Saccharomyces* and *Kluyveromyces* centromere/kinetochore complex proteins (named either Cep3(p) or CBF3 subunit B) with PnPf2 MHR (purple). Each protein structural superposition alignment shows the accompanying PDB accession and Cα RMSD value (as calculated by PyMOL CEAlign function).

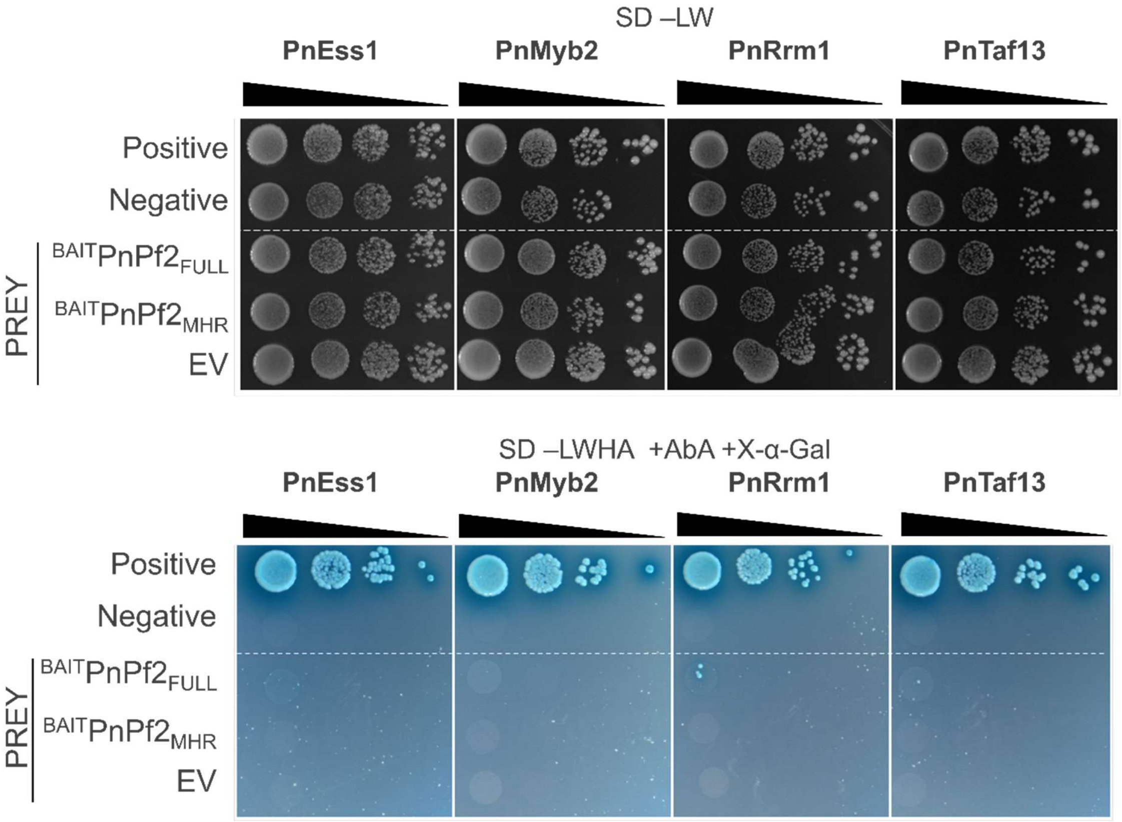


**Supplemental Figure S7**: Targeted Y2H of additional nuclear localising candidates against PnPf2. Additional targeted Y2H assays were performed for prey PnEss1, PnMyb2, PnRrm1 and PnTaf13 against PnPf2 full-length ($^{BAIT}PnPf2_{FULL}$) and PnPf2 MHR-domain only ($^{BAIT}PnPf2_{MHR}$). No significant growth was observed for any of tests, indicating these candidates did not interact with PnPf2 in a targeted Y2H assay.
